# Follicle cell contact maintains main body axis polarity in the *Drosophila melanogaster* oocyte

**DOI:** 10.1101/2022.03.03.482911

**Authors:** Ana Milas, Jorge de-Carvalho, Ivo A. Telley

## Abstract

In *Drosophila melanogaster* the anterior-posterior body axis is maternally established and governed by differential localization of partitioning defective (Par) proteins within the oocyte. At mid-oogenesis, Par-1 accumulates at the posterior end of the oocyte while Par-3/Bazooka is excluded there but maintains its localization along the remaining oocyte cortex. This mutual exclusion leads to a polarized microtubule network and accumulation of posterior determinant *oskar* later in oogenesis. Reciprocal biochemical interactions between Par proteins can explain their cortical exclusion and domain formation – for example, Par-1 excludes Par-3 by phosphorylation. However, past studies have proposed the need for somatic cells at the posterior end to initiate oocyte polarization by providing a trigger signal. To date, despite modern screening approaches and genetic manipulation, neither the molecular identity nor the nature of the signal is known. Here, we provide the first evidence that mechanical contact of posterior follicle cells (PFCs) with the oocyte cortex causes the posterior exclusion of Bazooka and maintains oocyte polarity. We show that Bazooka prematurely accumulates exclusively where posterior follicle cells have been mechanically detached or ablated. This occurs before Par-1 is removed suggesting that phosphorylation of Bazooka by Par-1 is not sufficient to maintain Bazooka exclusion in the absence of PFC contact. Furthermore, we provide evidence that PFC contact maintains Par-1 and *oskar* localization and microtubule cytoskeleton polarity in the oocyte. Our observations suggest that cell-cell contact mechanics modulates Par protein binding sites at the oocyte cortex.

## Introduction

A large majority of animals form two embryonic body axes (Niers, 2010). In many animals, these body axes are formed after fertilization, during early embryonic growth and segmentation. In *Drosophila melanogaster*, the anterior-posterior axis is maternally established, gradually during 14 stages of oogenesis, with the final goal of delivering *oskar* mRNA to the posterior end and *bicoid* mRNA to the anterior end of the oocyte (Riechmann and Ephrussi, 2001). The first sign of anterior-posterior polarity is the positioning of the oocyte to the posterior of the egg chamber at stage 1, which is achieved through differential adhesion between the oocyte and the somatic cells at the posterior of the egg chamber (Godt and Tepass, 1998; González-Reyes and St Johnston, 1998).

At stage 6, the posteriorly positioned oocyte secrets the EGF-like ligand, Gurken, which will cause the subset of follicle cells at the posterior to adopt posterior fate. In the following stage, these posterior follicle cells (PFCs) will signal back to the oocyte to determine its posterior pole (González-Reyes and St Johnston, 1994; González-Reyes et al., 1995; Roth et al., 1995). However, the molecular nature of the signal coming back from the PFCs remains elusive. Additionally, it is unclear if this signal acts solely as trigger that breaks symmetry, or if PFCs play a role in maintenance of oocyte polarity.

The earliest known signature of mid-stage oocyte polarization following the signal from PFCs is the di-phosphorylation of non-muscle Myosin II at the posterior of the oocyte (Doerflinger et al., 2022). This is necessary for posterior localization of Par-1 kinase, which is a component of a highly conserved network of Partitioning defective (Par) proteins (Shulman et al., 2000; Tomancak et al., 2000). Following Par-1 localization to the posterior, the anterior group of Par proteins, including aPKC, Par-6 and Par-3/Bazooka, relocalizes from the posterior to the anterolateral cortex (Doerflinger et al., 2010; Jouette et al., 2019). Posteriorly localized Par-1 inhibits nucleation of microtubules, causing plus ends of microtubules to preferentially accumulate at the posterior (Nashchekin et al., 2016; Parton et al., 2011). Polarization of the microtubule network is necessary to direct kinesin-dependent transport of *oskar* mRNA to the posterior during stages 9 and 10A (Lu et al., 2018, 2020; Zimyanin et al., 2008). Therefore, it is crucial that the polarity of the Par network is maintained during these stages.

The asymmetry of the Par network is thought to be self-maintaining through mutual antagonism between anterior and posterior Par proteins (Hoege and Hyman, 2013; Lang and Munro, 2017; Motegi and Seydoux, 2013). Par-1 phosphorylates Bazooka to exclude the complex of aPKC/Par-6/Bazooka from the posterior, while aPKC phosphorylates Par-1 leading to the removal of Par-1 from the anterolateral membrane (Benton and St Johnston, 2003a; Doerflinger et al., 2010). Interestingly, anterior and posterior Par proteins colocalize at the posterior of the oocyte from stage 7 to stage 9, suggesting that mutual antagonism might not be sufficient to maintain Par polarity (Doerflinger et al., 2010; Jouette et al., 2019).

Here, we first show that at late stage 10B of oogenesis the cell-cell interface of the oocyte and PFCs visibly enlarges, suggesting contact loss, followed by accumulation of Bazooka and removal of Par-1 at the posterior of the oocyte. Mechanical detachment of PFCs from the oocyte at stage 9 and 10A by micromanipulation causes loss of contact and premature accumulation of Bazooka. The exclusion of Bazooka from the posterior is rapidly reverted following laser ablation of PFCs. Bazooka accumulates at the posterior before Par-1 is removed, suggesting that Par-1 phosphorylation of Bazooka is not sufficient to maintain Bazooka exclusion once PFC contact is lost. Following PFC ablation, Par-1 delocalizes slowly leading to loss of *oskar* mRNA localization at the posterior, but this occurs faster than the rate at which Par-1 targeting kinase aPKC accumulates. Finally, microtubules polymerize at the posterior, but after *oskar* delocalizes. We conclude that PFCs maintain polarity of the Par network by cell-cell contact until late stage 10B of oogenesis by excluding Bazooka from the posterior of the oocyte. This is necessary to anchor *oskar* mRNA at the posterior, and to maintain polarization of the microtubule network during stages at which *oskar* is delivered to the posterior by directed transport.

## Results

### Posterior relocalization of Bazooka following contact loss with PFCs in stage 10B oocytes

The localization patterns of Bazooka have been well described up until early stage 10 of oogenesis. At stages 7 and 8, Bazooka localizes to the posterior of the oocyte, where it overlaps with the Par-1 domain, until stage 9 when it is excluded from the posterior (Doerflinger et al., 2010; Jouette et al., 2019). However, the dynamics of Bazooka localization in the following stages is not known. To assess this, we performed live cell imaging of stage 10 and 11 egg chambers expressing Bazooka tagged with GFP (Baz::GFP) and Jupiter tagged with mCherry (Jupiter::mCherry) as a reporter for microtubules (Fig. 1). We used the GAL4/UASp system to express Bazooka only in the germline, since Bazooka localizes to the apical side of follicle cells, thus masking the localization of Bazooka at the oocyte membrane (Jouette et al., 2019). In agreement with previous results, all but one oocyte show exclusion of Bazooka at stage 10A (n=13). More interestingly, the oocyte is tightly connected to the PFCs at this stage, i.e., a clear cellular boundary between the oocyte and the PFCs is not detected in fluorescence confocal images of microtubules and in polarized transmission light microscopy images (Suppl. Fig. 1A, arrowheads). On the other hand, there is a visible gap between the lateral follicle cells and the oocyte. This differential contact correlated with the accumulation of Bazooka; Bazooka localizes at the membrane where the oocyte is not in contact with the follicle cells and is excluded where the contact is established (Suppl. Fig. 1A). At stage 11, the cell-cell contact between the oocyte and the PFCs is lost, and a clear boundary is visible, while the exclusion of Bazooka is lost as well (Suppl. Fig. 1B, arrowhead). To understand if the formation of an intercellular space (gap) between PFCs and the oocytes precedes or follows the accumulation of Bazooka at the posterior, we analyzed stage 10B egg chambers in which this transition likely occurs (Fig. 1A–C). The posterior gap is visible in 83% (30 out of 36) stage 10B egg chambers. However, Bazooka was still excluded from the posterior in 12 of these oocytes (Fig. 1D). Importantly, we did not observe egg chambers where Bazooka accumulated to the posterior before the gap formed. The inter-cellular gap forms first anterolaterally sometime during stage 9, before Bazooka is excluded (Fig. 1D). We conclude that the loss of contact between the oocyte and PFCs precedes accumulation of Bazooka (Fig. 1E).

**Figure 1.**
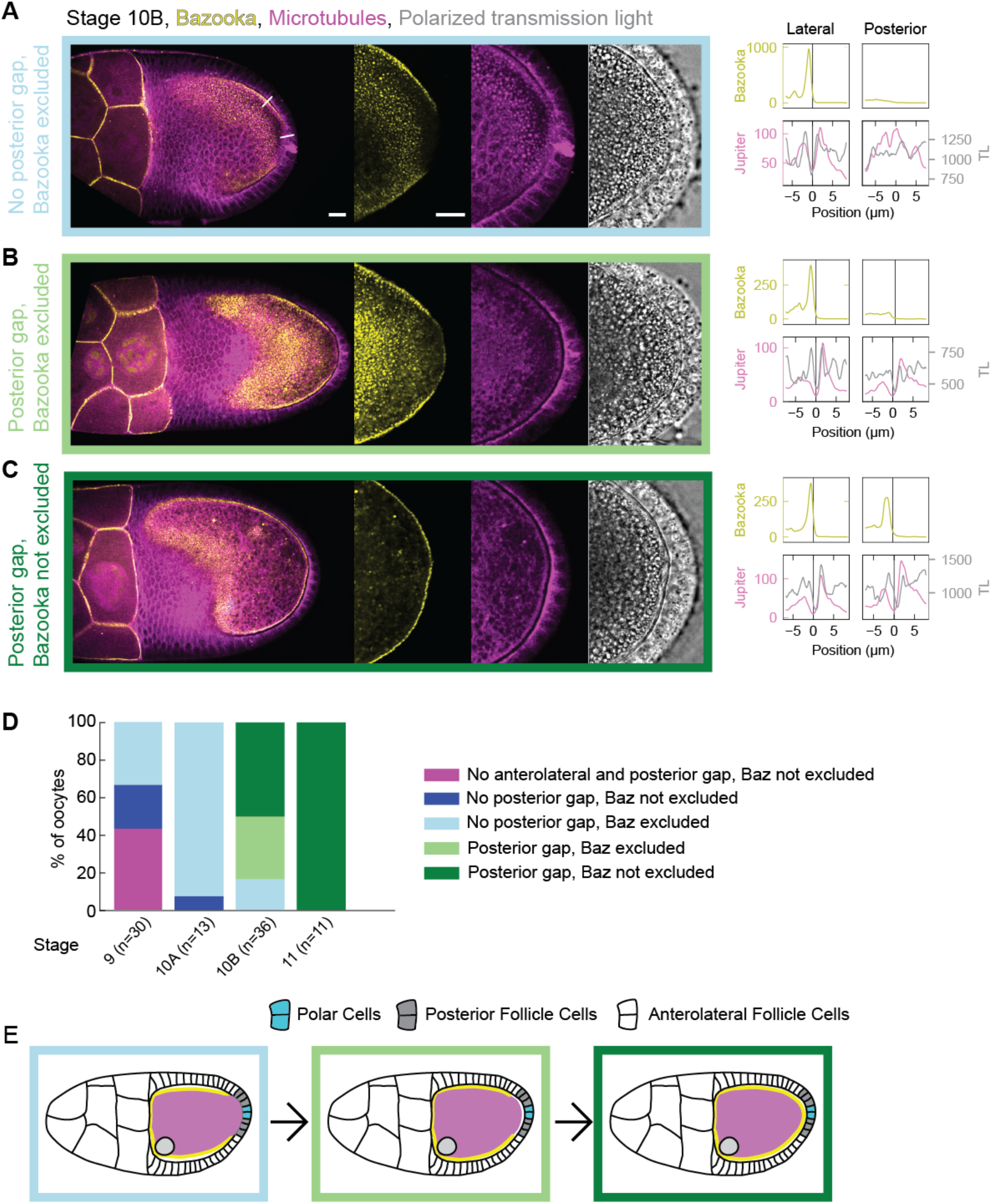
Bazooka accumulates at the posterior following the loss of contact between the oocyte and PFCs at stage 10B of oogenesis: **(A–C)** On the left are still images of stage 10B egg chambers expressing Bazooka::GFP (yellow) and the microtubule reporter Jupiter::mCherry (magenta) next to a transmission light micrograph (grey). The right graphs are intensity profiles of Bazooka::GFP (yellow), Jupiter::mCherry (magenta) and transmission light (grey) signal along a straight line crossing cell boundaries from the oocyte to either lateral or posterior follicle cells (see first image of the panel A). The x-axis origin and vertical line represent the local minimum of the Jupiter::mCherry (magenta) signal, which we interpret as extended intercellular space (gap) between the oocyte and the follicle cells. Whenever this intercellular space is not clearly discernible, the position 0 μm represents the midpoint of the line. **(A)** At the beginning of stage 10B, a gap is detected between the oocyte and the lateral follicle cells but not the posterior follicle cells. Accumulation of Bazooka at the oocyte membrane correlates with existence of the gap. **(B)** Later in stage 10B, a gap is formed at the posterior, but Bazooka does not yet accumulate at the respective oocyte membrane. **(C)** Eventually, Bazooka accumulates to the posterior after the formation of the gap. The scale bars represent 20 μm for panels A–C. **(D)** Distribution of egg chambers showing one of the four possible phenotypes at stages 9, 10A, 10B and 11. Stage 9 is distinct from later stages in that a fraction of egg chambers shows no (anterolateral) gap at all. The colors in the plot correspond to the color of the frame surrounding the images in panels A–C. *n* is the number of egg chambers quantified. **(E)** Scheme showing the timeline of events during stage 10B. Loss of contact between the oocyte and PFCs precedes accumulation of Bazooka to the posterior.

### Bazooka accumulates at the posterior before Par-1 clearance at late stage 10B of oogenesis

In *par1* mutant oocytes Bazooka localizes all around the cortex, suggesting that Par-1 phosphorylates Bazooka to exclude it from the posterior (Benton and St Johnston, 2003a). To assess if Par-1 needs to be removed from the posterior before Bazooka accumulates, we imaged oocytes expressing fluorescently tagged Par-1 and Bazooka. At stage 10A, we found Bazooka exclusion and Par-1 enrichment in all analyzed oocytes (n=12) (Suppl. Fig. 1C–F). On the other hand, at stage 11, differential localization of both Bazooka and Par-1 at the posterior were lost (n=10) (Suppl. Fig. 1E–F). At stage 10B, Bazooka relocalized to the posterior in 25 out of 40 oocytes. However, Par-1 was still present at the posterior in 12 of these oocytes (Suppl. Fig. 1D, F). Importantly, we never found Par-1 disappearing from the posterior before accumulation of Bazooka. Therefore, Bazooka first accumulates to the posterior, which eventually leads to the removal of Par-1 from this region. This result suggests that Bazooka accumulation is not the consequence of Par-1 removal, and that Par-1 is not sufficient to exclude Bazooka from the posterior during this transition.

Taken together, these observations suggest that the loss of contact between PFCs and the oocyte is followed first by recruitment of Bazooka to the posterior, and only thereafter Par-1 delocalizes from the posterior. We hypothesize that cell-cell contact between PFCs and the oocyte is required to exclude Bazooka from the posterior oocyte membrane.

### Mechanical detachment of PFCs from the oocyte causes posterior accumulation of Bazooka

To directly test the contact interaction between oocyte and PFCs leading to the exclusion of Bazooka, we designed an experiment by which PFCs were mechanically detached from the oocyte. We used a blunt glass micropipette mounted on a micromanipulator to aspirate and pull on the PFCs aiming detachment from the oocyte but keeping them intact (Fig. 2A). We combined this assay with live imaging of egg chambers expressing Baz::GFP in the germline and Jupiter::mCherry in all tissues to observe any changes in polarity in the oocyte (Fig. 2B and Video 1). Indeed, we could observe accumulation of Bazooka to the posterior of the oocyte following detachment of the PFCs (Fig. 2B–C). Importantly, this accumulation was accompanied by the appearance of the intercellular space between oocyte and PFC boundaries (Fig. 2D). We conclude that mechanical pulling on PFCs causes premature intercellular gap formation, which results in the accumulation of Bazooka to the posterior of the oocyte. This suggests that a firm cell-cell interaction between PFCs and the oocyte is important for maintaining Bazooka exclusion at the posterior end of the oocyte.

**Figure 2.**
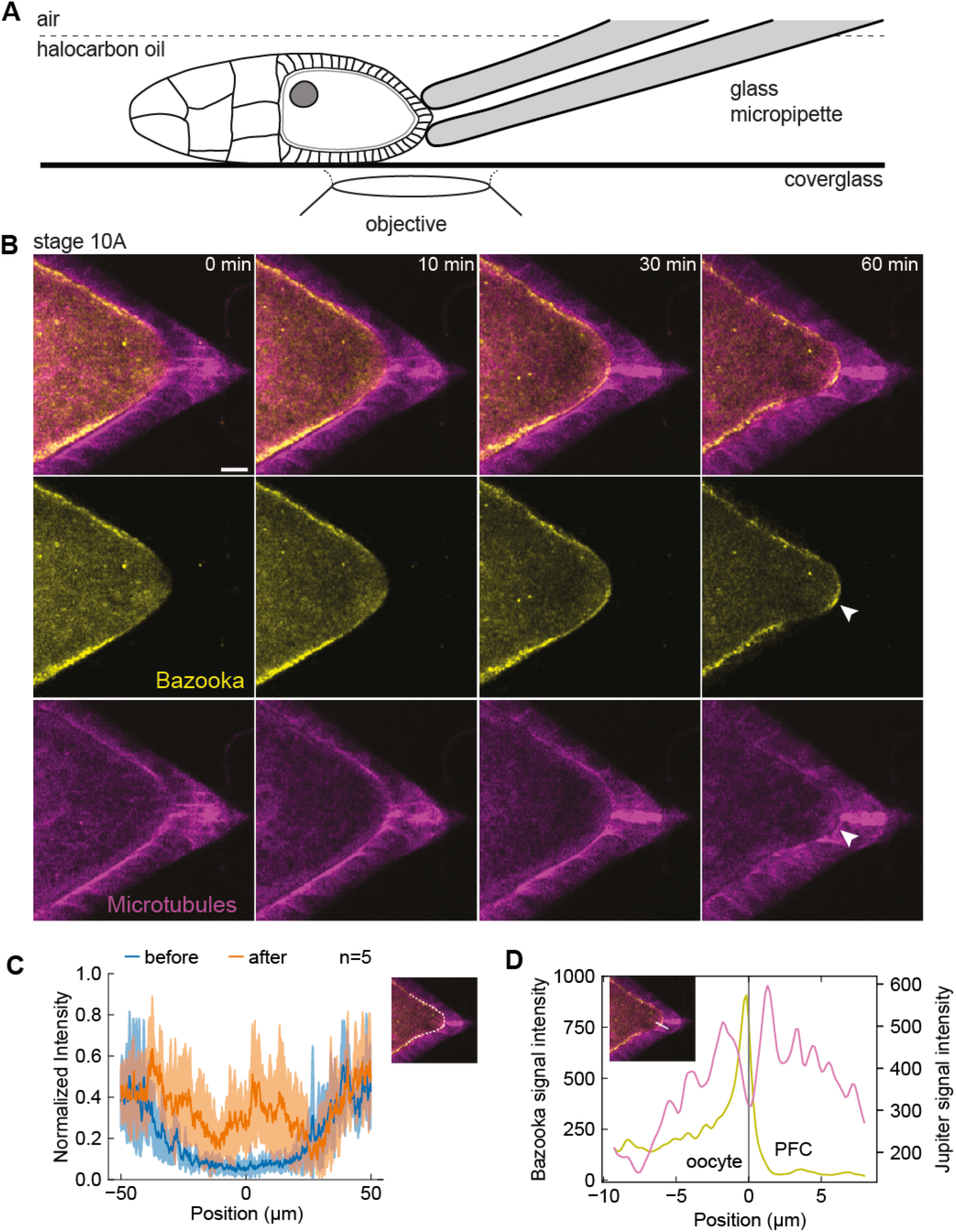
Mechanical detachment of PFCs from the oocyte causes posterior accumulation of Bazooka: **(A)** Schematic showing experimental assay to mechanically detach PFCs from the oocyte. A holding micropipette is used to aspirate and pull on the PFCs. **(B)** Time-lapse images of an egg chamber expressing Bazooka::GFP (yellow) and the microtubule reporter Jupiter::mCherry (magenta) following detachment of PFCs from the oocyte. Top: merged channels, middle: Bazooka::GFP, bottom: Jupiter::mCherry. Formation of the intercellular space and accumulation of Bazooka are detected at the posterior (arrowheads). Scale bar, 20 μm **(C)** The average intensity profile of Bazooka::GFP signal at the posterior of the oocyte, measured along the oocyte cortex as represented by the white line in the inset image. The blue curve is the intensity measured before the PFCs were pulled, the orange curve is the intensity after 60 min of continuous pulling. The shaded region designates the standard deviation. The x-axis origin represents the position of the polar cells. *n* is the number of experiments. **(D)** Intensity profile of Bazooka::GFP (yellow) and Jupiter::mCherry (magenta) along the line crossing cell boundaries from the oocyte to posterior follicle cells, as shown in the inset. The x-axis origin and vertical line represent the local minimum of the Jupiter::mCherry signal, which we interpret as intercellular space between the oocyte and the follicle cells.

### Posterior follicle cells maintain posterior exclusion of Bazooka with cell size-precision

Since cell differentiation of follicle cells into PFCs is necessary to establish polarity of the oocyte in the first place, mutants that disrupt their differentiation do not allow testing of their role in polarity maintenance throughout oogenesis. Therefore, we sought of a more acute and spatially targeted perturbation method for disrupting PFCs. We used pulsed UV laser ablation to destroy a small number of PFCs and monitor the resulting changes in Par protein localization in the oocyte, comparing locations where PFCs were removed versus where they are intact. First, we performed ablation in stage 9 or 10A egg chambers that express Baz::GFP in the germline and Jupiter::mCherry in all tissues. In every experiment, Bazooka was excluded from the posterior prior to the ablation (Fig. 3A, t = −1 min). The ablation of a few PFCs resulted in accumulation of Bazooka exclusively to the membrane region facing the ablated cells (Fig. 3B–C and Video 2). Bazooka did not localize at neighboring regions where PFCs were intact (Fig. 3B, arrowheads). A quantification of Baz::GFP intensity along the posterior cortex of the oocyte showed a signal peak confined to the location of ablated PFCs (Fig. 3C, vertical lines). Similar results were obtained when ablated posterior follicle cells did not include the pair of polar cells (Suppl. Fig. 2A–C, Fig. 1E). To account for possible turnover kinetics and photobleaching, we compared Baz::GFP intensity at cortices either facing posterior or facing lateral follicle cells (Fig. 3D, inset), confirming a significant increase and in some experiments a full recovery of Baz::GFP localization at the posterior. A temporal analysis of Baz::GFP intensity after PFC ablation revealed an almost immediate increase (∼1 min delay) (Fig. 3E). The accumulation seemed faster and achieved steady state earlier if polar cells were left intact. We generated a spatiotemporal map of average Baz::GFP intensity along the oocyte cortex facing ablated PFCs (Fig. 3F–G). Whenever the ablated region contained the pair of polar cells, Bazooka accumulated in random patches, presumably by being recruited from the cytoplasmic pool (Fig. 3F). In contrast, when the ablated region was flanked by still intact polar cells on one side, and main body follicle cells on the other side, we noticed a signal flow from the anterolateral cortex facing main body follicle cells. However, we did not notice any significant cytoplasmic flow in the time-lapse movies. We concluded that Bazooka predominantly diffused along the membrane from the cortical pool of anterolateral side towards the posterior end.

**Figure 3.**
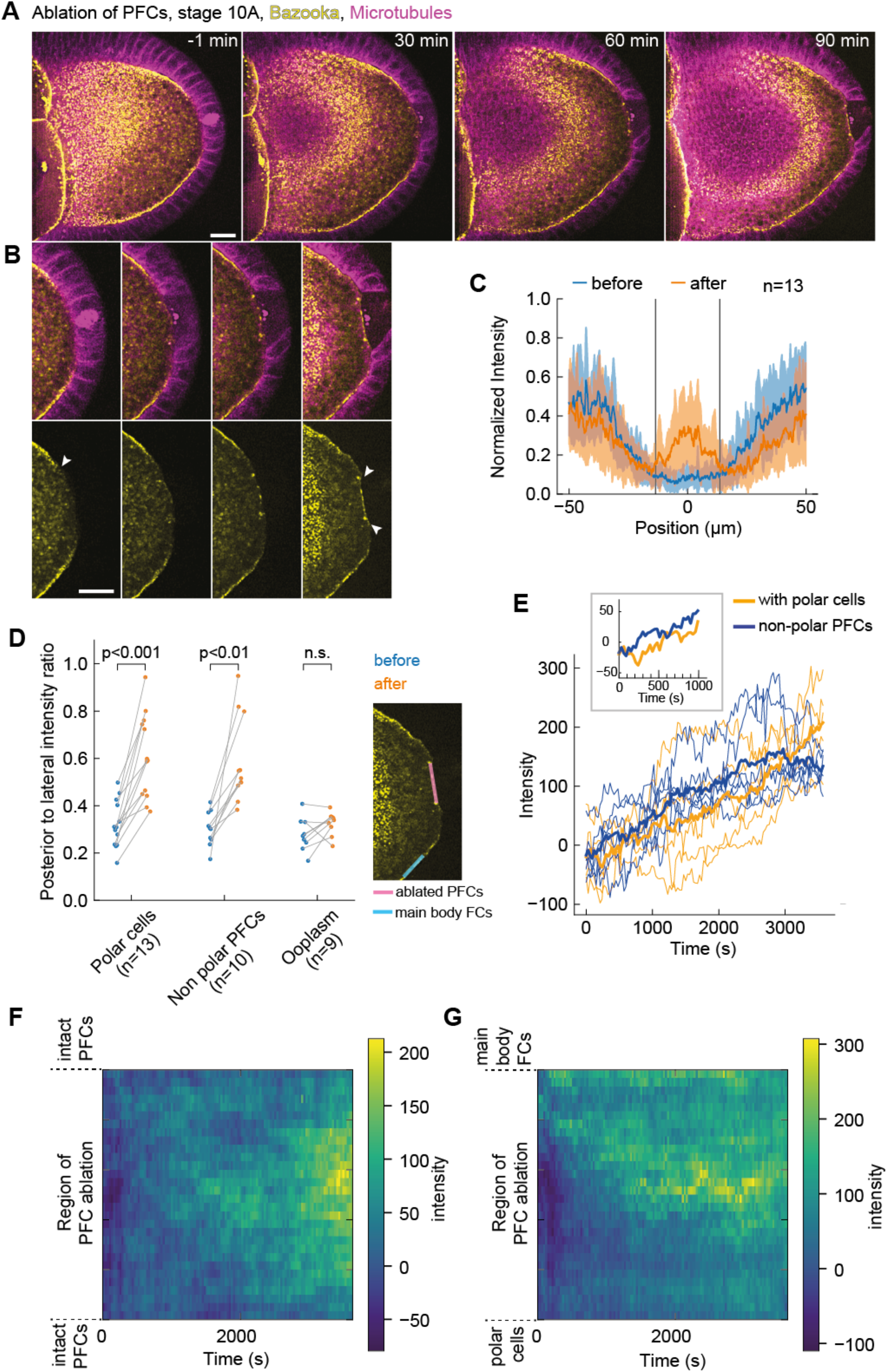
Bazooka localizes to the posterior following ablation of PFCs: **(A)** Two-color time-lapse images of an egg chamber expressing Bazooka::GFP in the germline (yellow) and the microtubule reporter Jupiter::mCherry (magenta) before (-1 min) and after ablation of PFCs. Scale bar, 20 μm. **(B)** Zoom-in of the posterior end of the egg chamber shown in panel A. Top: merged channels, bottom: Bazooka::GFP. Before ablation, Bazooka is excluded from the posterior, a region highlighted by the arrowheads in the first image of the bottom panel. After 90 min, accumulation of Bazooka is visible at the oocyte membrane facing the ablated cells (arrowheads in the last image). Scale bar, 20 μm. **(C)** Average intensity profile of Bazooka::GFP signal at the posterior of the oocyte before (blue) and after (orange) ablation. The shaded region designates the standard deviation. The x-axis origin represents the position of the ablated region. Vertical lines denote the average width of the ablated region. Note that the increase of the signal is visible only inside the ablated region. **(D)** Ratio between average Bazooka::GFP intensities along the oocyte membrane facing ablated PFCs and lateral main body follicle cells, as depicted in the inset image, before (blue) and after (orange) ablation for three types of ablation. PFCs: ablation of PFCs, including ablation of polar cells (as in panel A). Non-polar PFCs: ablation of PFCs that do not include polar cells (see Suppl. Fig. 2A). Ooplasm: ablation inside the ooplasm (see Suppl. Fig 2D). A ratio equal to 1 means complete loss of Bazooka exclusion. Wilcoxon signed rank test was performed to assess significance (n.s.=not significant). **(E)** Intensity of Bazooka::GFP signal at the oocyte membrane facing the ablated follicle cells as a function of time, grouped in experiments where polar cells were ablated (orange, n=5) or left intact (blue, n=5). The thick lines represent the average of each group, thin colored lines are individual oocytes. Intensity of the ooplasm was subtracted from the Bazooka::GFP intensity, so that zero intensity (on average) means exclusion of Bazooka. Inset: Average curves in the first 15 min. **(F)** Time series of Bazooka::GFP intensity along the oocyte membrane facing ablated PFCs (y-axis) after PFC ablation, represented as heat map. The ablated follicle cells included the polar cells, and the ablated region is flanked by still intact PFCs. **(G)** As in panel F, but for ablations that excluded polar cells, so that the ablated region was flanked by intact polar cells on one side (bottom of y-axis) and main body follicle cells on the other side (top of y-axis). Note the signal inflow from the top of the graph. For all panels t = 0 is the first frame after ablation, *n* is the number of experiments.

To assure that accumulation of Bazooka is not caused by nonspecific effects of pulsed laser ablation, we performed ablation inside the ooplasm, close to the posterior membrane of the oocyte (Suppl. Fig. 2D and Video 3). We did not observe any accumulation of Bazooka at the oocyte membrane, or around the region that was ablated (Suppl. Fig. 3E–F). We also addressed if ablation of follicle cells may cause nonspecific accumulation of any protein to the membrane. Thus, as additional control experiment, we UV targeted main body (anterolateral) follicle cells in egg chambers expressing Par-1::GFP in the germline and Jupiter::mCherry. At this stage, Par-1 is restricted to the posterior of the oocyte and there is no Par-1 signal at the oocyte membrane in contact with main body follicle cells (Suppl. Fig. 3A, left). Importantly, we did not observe any increase in GFP signal following ablation (Suppl. Fig. 3B–C). From all these experiments, we conclude that individual PFCs are required to locally maintain posterior exclusion of Bazooka throughout mid-oogenesis.

### Stage dependent maintenance of Par-1 localization by PFCs

Next, we wanted to test if Par-1 needs to be excluded from the posterior before Bazooka can accumulate, upon ablation of PFCs. According to the mutual exclusion hypothesis of the Par protein network, Par-1 presence is responsible for exclusion of the anterior Par protein complex, including Bazooka, through kinase activity (Benton and St Johnston, 2003a). Therefore, we performed the ablation experiments in egg chambers expressing Par-1::GFP and Baz::mCherry in the germline. With this combination of polarity protein reporters, we again observed accumulation of Bazooka to the posterior following the ablation of PFCs (Fig. 4A, Suppl. Fig. 3D and Video 4 and 5). A first analysis suggested that Par-1 does not decrease as rapidly as expected. Ectopic localization of Bazooka was preceded by a rather slow disappearance of Par-1. However, we suspected that the developmental stage marks a difference in Par-1 kinetics upon PFC ablation. In stage 10A oocytes, Par-1 signal marginally decreases within one hour (Suppl. Fig 3F) although longer acquisitions revealed a decrease. In stage 9 oocytes, the signal decrease was more pronounced in the region that had been in contact with ablated PFCs (Fig. 4B–C). Slow signal changes can be either a consequence of late response to the perturbation, or due to photobleaching. Thus, we performed a comparative analysis within each oocyte, and acquired at low light exposure (Methods). Ectopic accumulation of Bazooka at the posterior was significant when compared to the lateral cortex (Fig. 4D), while the Par-1 signal decrease was not significant in stage 10A (Fig. 4E). We conclude that Par-1 is not sufficient to keep Bazooka excluded from the posterior after PFC ablation. Moreover, Par-1 localization is controlled by PFCs in a stage dependent manner.

**Figure 4.**
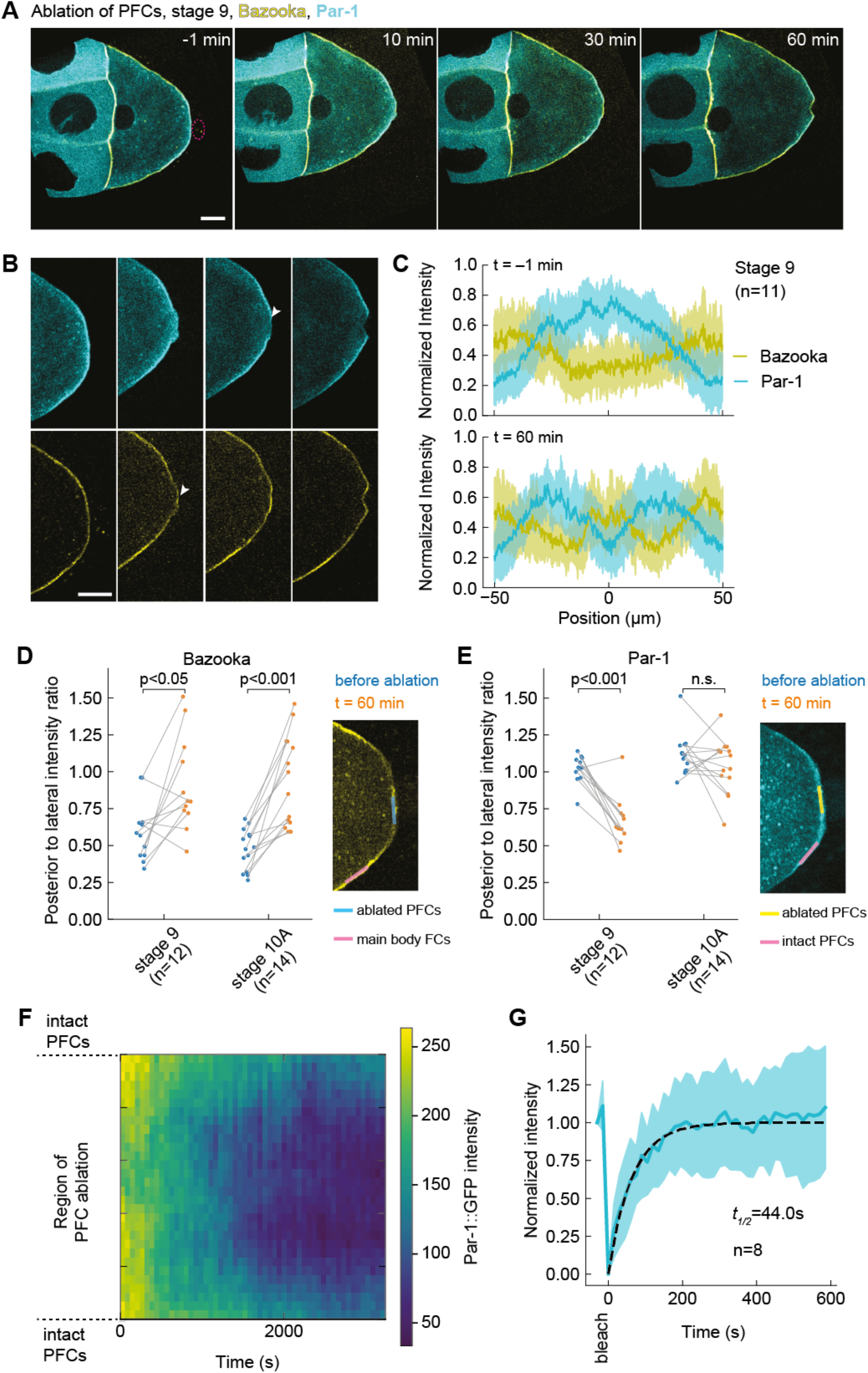
Par-1 delocalizes slowly from the posterior despite fast turnover: **(A)** Two-color time-lapse images of an egg chamber at stage 9 expressing Par1::GFP (cyan) and Bazooka::mCherry (yellow) in the germline, before (-1 min) and after ablation of PFCs (red dashed circle). Scale bar, 20 μm **(B)** Zoom into the posterior of the egg chamber shown in panel A. Top: Par1::GFP, bottom: Bazooka::mCherry. Following ablation, Bazooka accumulates to the posterior, while Par-1 is still present but slowly delocalizing (arrowheads). Scale bar, 20 μm. **(C)** Average intensity profile of Par-1::GFP (cyan) and Bazooka::mCherry (yellow) signal at the posterior of stage 9 oocytes before (top graph) and one hour after ablation of PFCs (bottom graph). The shaded region designates the standard deviation. The x-axis origin represents the position of the polar cells (top) or center of ablation spot (bottom). **(D)** Ratio between Bazooka::mCherry intensities at the posterior and lateral membrane of the oocyte (facing main body follicle cells), as depicted in the inset image, before (blue) and one hour after (orange) ablation. The ratio 0 signifies complete posterior exclusion, 1 and beyond denote localization. There is a significant increase in the ratio for both stages 9 and 10A, showing accumulation of Bazooka to the posterior. **(E)** Ratio between Par-1::GFP intensities at the membrane of the oocyte facing posterior ablated PFCs and intact PFCs sightly lateral, as depicted in the inset image, before (blue) and one hour after (orange) ablation. There is no significant reduction of the ratio in stage 10A which would indicate delocalization of Par-1. Wilcoxon signed rank test was performed to assess significance (n.s. = not significant). **(F)** Time series of Par-1::GFP intensity along the oocyte membrane facing ablated PFCs (y-axis) after PFC ablation, represented as heat map. **(G)** Normalized signal recovery after fluorescence photobleaching for Par1::GFP signal at the oocyte membrane. Thick blue line is mean, the shaded region designates the standard deviation, the dashed line is the fitted curve obtained by fitting a single exponential function. For all panels, *n* is the number of experiments.

We noticed that Par-1 loss seemed to start in the center of the cortex region which lost contact with PFCs (Suppl. Fig. 3E, arrowhead), and extended towards the cortical boundaries facing intact PFCs. To obtain better insight, we generated a spatiotemporal map of Par-1::GFP intensity along the oocyte cortex facing the ablated PFCs (Fig. 4F). This analysis confirmed a progressive signal loss that occurs symmetrically around the ablation center. In an effort to decompose cortical mobility and turnover kinetics of Par-1::GFP at the posterior cortex, we performed FRAP analysis in the absence of perturbation. Typical Par-1::GFP recovery time was <1 min (Fig. 4G), which contrasts the slow Par-1 delocalization after perturbation, occurring typically within one hour (Fig. 4F). We did not notice any cytoplasmic flow which could potentially transport Par-1 to the boundaries by advection (Video 4 and 5). Thus, our interpretation is that the molecular binding sites for Par-1, which are abundant and cause Par-1 accumulation, slowly delocalize upon PFC ablation while Par-1 turns over fast. Importantly, the spatial signature of delocalization is neither uniform nor random; having excluded advection, it is reminiscent of diffusible binding sites whose mobility within the membrane is reduced when the cortex is in contact with PFCs (“modulated molecular trap”).

### Posterior *oskar* mRNA is lost upon PFC ablation

A downstream process of Par polarity in the oocyte is *oskar* mRNA accumulation at the posterior, which ultimately defines the tail of the future organism (Riechmann and Ephrussi, 2001). This is achieved by directed transport on a slightly polarized microtubule network and anchoring at the posterior cortex (Lu et al., 2018, 2020; Zimyanin et al., 2008). We wanted to understand if the response of Par proteins to PFC ablation would also affect *oskar* localization. To this end, we performed live cell imaging of stage 8–10A egg chambers expressing Par-1::GFP and *osk*MS2 under its native driver together with MCP::mCherry (Fig. 5A, Video 6). Upon PFC ablation, Par-1 delocalizes slowly, as shown previously (Fig. 5B, top). More strikingly, *oskar* also delocalizes, first in the region previously in contact with ablated PFCs, and gradually also at the flanking regions (Fig. 5B, bottom). Qualitatively, *oskar* delocalization seemed delayed and slower for later stages. An analysis of the signal kinetics for both Par-1 and *oskar* by fitting single exponential decay functions with delay confirmed this impression (Fig. 5C). *oskar* delocalization occurred with a statistically significant delay relative to Par-1. In stage 8 oocytes the overall delocalization kinetics was in the minutes range, while in stage 10A, in particular for *oskar*, the delay and kinetics were in the hour range. We conclude that PFCs are required to maintain body axis definition in the oocyte until stage 10 by defining *oskar* mRNA localization through Par-1.

**Figure 5.**
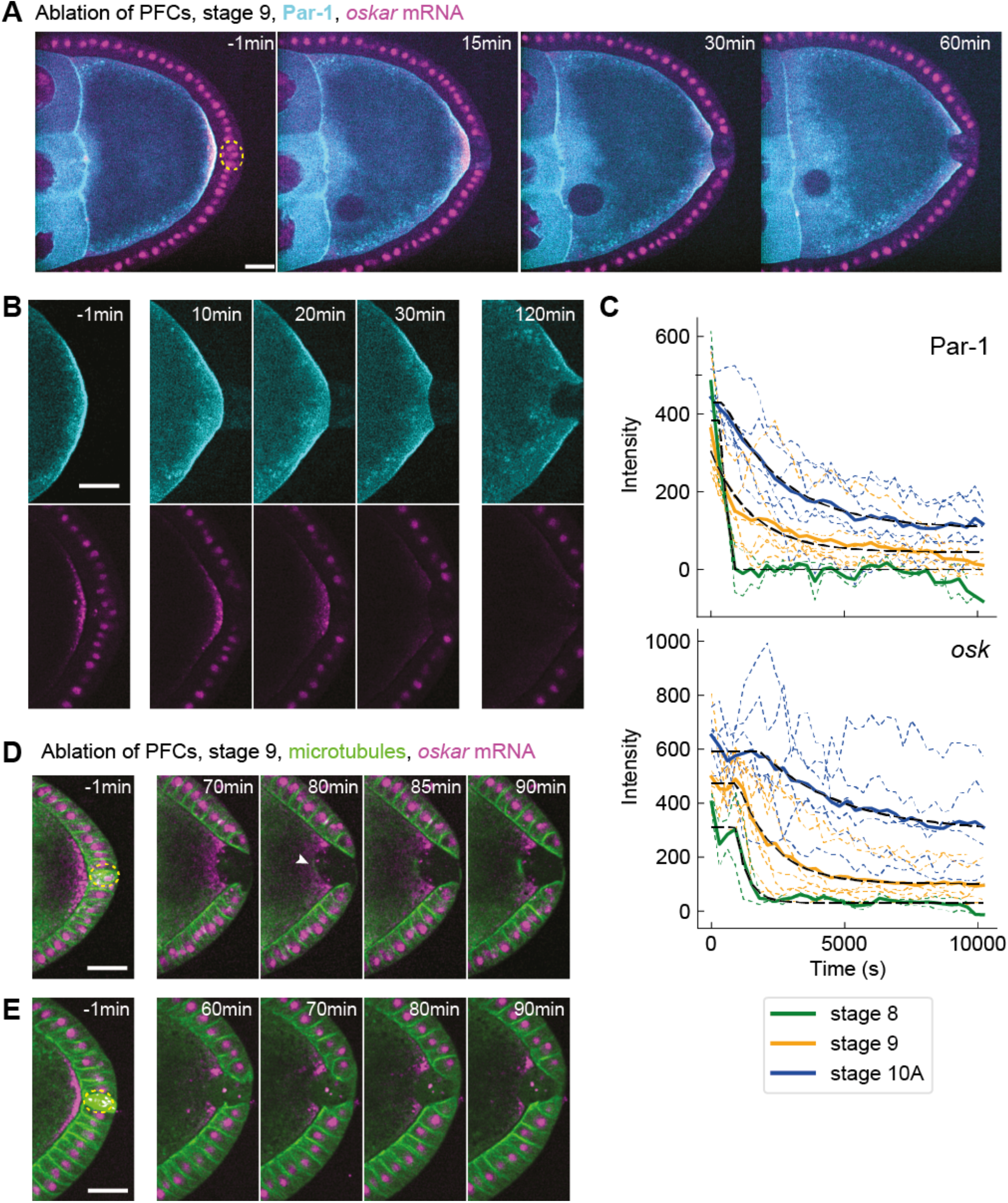
Posterior *oskar* mRNA is lost shortly after Par-1, and microtubules polymerize with long delay: **(A)** Two-color time-lapse images of an egg chamber expressing Par1::GFP in the germline (cyan) and *oskMS2*-mCherry (magenta) before (-1 min) and after ablation of PFCs (yellow dashed circle). Scale bar, 20 μm **(B)** Zoom-in of the posterior of the egg chamber shown in panel A. Top: Par-1::GFP, bottom: *oskMS2*-MCP::mCherry. Both Par-1 and *osk* delocalize from the cortex facing ablated PFCs. Scale bar, 20 μm. **(C)** Intensity of Par-1 (top) and *osk* mRNA (bottom) at the posterior of the oocyte facing ablated PFCs, for stages 8 (n=2), 9 (n=7) and 10A (n=6). Thin dashed lines are individual experiments, the solid lines are the averages for each stage, and the black dashed line represents a fit to a single exponential decay with time delay (see Methods). While Par-1 signal loss is immediate, *osk* mRNA decreases with increasing delay; the estimated 95% CI (min) are for stage 8: [13.3, 16.5], for stage 9: [15.5, 18.9], for stage 10A: [24.4, 36.0]. **(D)** Two-color time-lapse images of an egg chamber expressing Jupiter::GFP (green) and *oskMS2*-mCherry (magenta) before (-1 min) and after ablation of PFCs. This example shows microtubule growth at the posterior 80–90 min following ablation. Scale bar, 20 μm. **(E)** As in panel D, showing an example of an oocyte that did not yet exhibit microtubule growth 90 min after PFC ablation. Scale bar, 20 μm.

Finally, motivated by earlier reports of Par-1 being a microtubule nucleation inhibitor (Doerflinger et al., 2010; Nashchekin et al., 2016; Parton et al., 2011), we predicted that the delocalization of Par-1 upon PFC ablation should enable microtubule growth in that region. Growth of microtubules from the posterior towards the cytoplasm could explain the loss of *oskar* due to directed transport away from the cortex. Indeed, in some oocytes we observed a local signal increase of the microtubule reporter Jupiter::GFP where *oskar* signal decreased (Fig. 5D and Video 7). However, growth initiated considerably later than *oskar* delocalization, so that in some oocytes we did not observe any growth within the time of observation (Fig. 5E). Thus, this result shows that the maintenance of Par polarity by PFCs is functionally important for robust microtubule cytoskeleton polarization. However, loss of *oskar* upon PFC ablation is likely not caused by microtubule reorganization but rather from loss of cortical anchoring.

### aPKC kinetics after PFC ablation does not explain Par-1 removal from the posterior

According to the current model of Par network polarization in the *Drosophila* oocyte, Par-1 is excluded from the anterolateral membrane through phosphorylation by the kinase aPKC, which is recruited to the membrane through interaction with Bazooka (Doerflinger et al., 2010). Conceivably, accumulation of Bazooka at the posterior after PFC ablation may recruit aPKC, eventually leading to Par-1 delocalization at the posterior. To obtain insight into the sequence of events, we performed live imaging of egg chambers from flies that endogenously express aPKC::GFP (Fig. 6A and Video 8) (Chen et al., 2018) and Bazooka::mCherry in the germline. Since both follicle cells and germline express aPKC, the initial posterior exclusion of aPKC in the oocyte is masked by the apical signal in follicle cells. However, once PFCs are ablated their aPKC::GFP signal disappears and any fluorescence detected can be associated to aPKC in the oocyte (Fig. 6B, bottom row). Therefore, our assay allows us to unambiguously detect possible accumulation of aPKC following ablation of PFCs. Although the signal of endogenously driven protein expression is significantly lower when compared to using GAL4/UASp expression system, we were able to observe accumulation of aPKC in most of the oocytes (Fig. 6B, arrowheads). An intensity analysis of aPKC::GFP showed significant accumulation at stage 9. More importantly, aPKC localization lagged upon perturbation and was always preceded by the accumulation of Bazooka (Fig. 6C–D). Interestingly, we did not detect any significant increase in aPKC at stage 10A (Fig. 6D), which could be explained by the signal detection limit in our imaging. More assuringly, the loss of Par-1 was on average faster than the kinetics of aPKC ectopic localization after perturbation (Fig. 6E), suggesting that Par-1 loss occurred before aPKC localized. We conclude that aPKC accumulation is insufficient for Par-1 displacement, and the maintenance of Par-1 localization at the posterior requires the oocyte to be in contact with PFCs.

**Figure 6.**
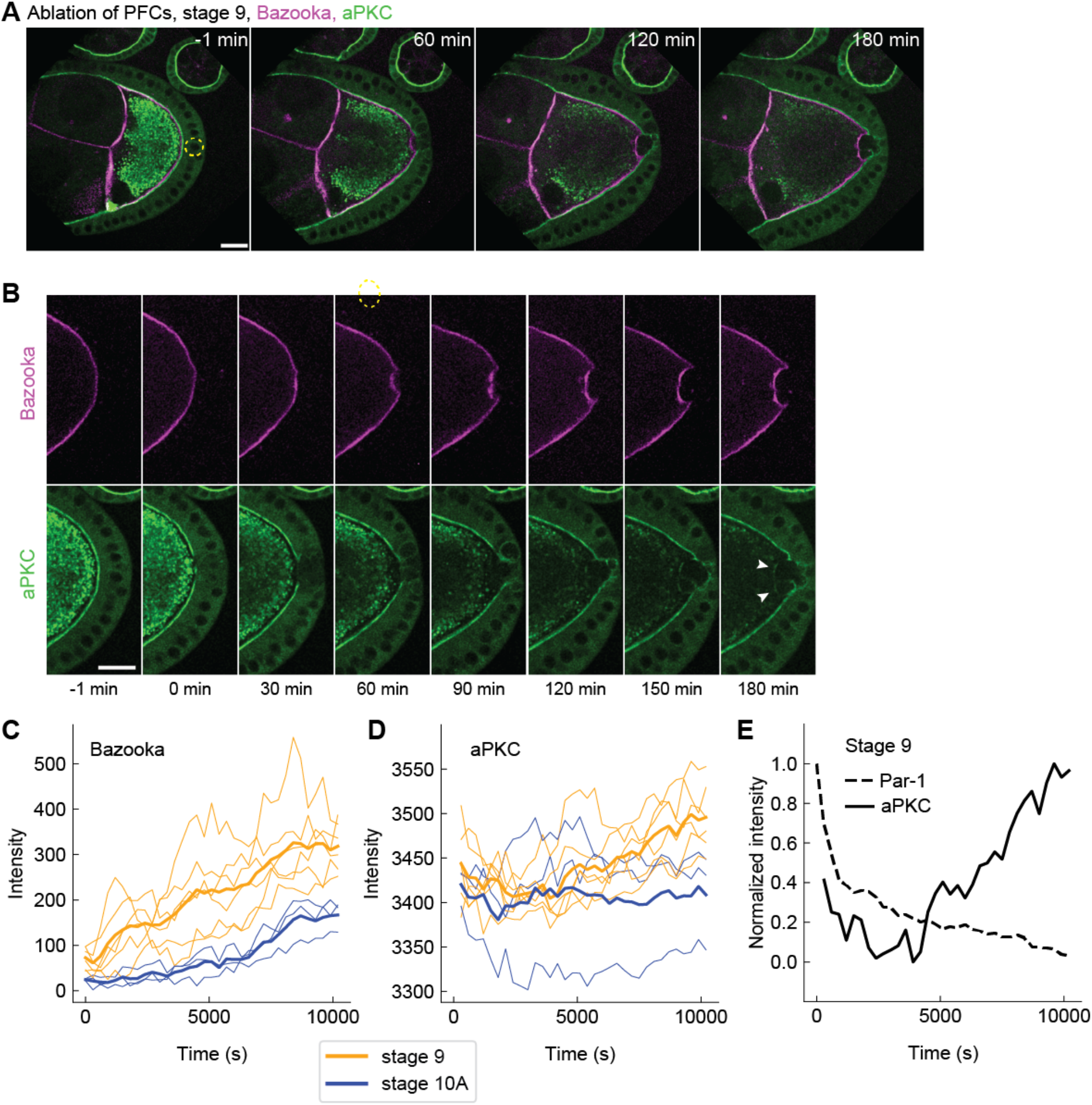
aPKC accumulates at the posterior after the changes in Bazooka and Par-1. **(A)** Two-color time-lapse images of an egg chamber at stage 9, expressing endogenous aPKC tagged with GFP (green) and Bazooka-mCherry in the germline (magenta) before (-1 min) and after ablation of PFCs (yellow dashed circle). Scale bar, 20 μm **(B)** Zoom-in of the posterior of the egg chamber shown in panel A. Top: Bazooka::mCherry, bottom: aPKC::GFP. While Bazooka accumulates within 60 minutes, aPKC is detectable only after 120–150 min (arrowheads). Scale bar, 20 μm. **(C)** Bazooka::mCherry intensity at the oocyte cortex facing the ablated PFCs, as a function of time for oocytes at stage 9 (orange, n=6) and 10A (blue, n=3). Thin lines are individual experiments, thick lines are stage averages. **(D)** As in panel C but for aPKC::GFP. Note the delayed increase in stage 9, and the lack of signal increase in stage 10A. **(E)** Normalized average intensities of Par1::GFP (dashed line) and aPKC::GFP (solid line) at stage 9 as a function of time following the ablation of PFCs. Note the delay of aPKC signal increase. For all panels, time zero marks the first frame after ablation.

## Discussion

In *Drosophila*, the canonical events of oocyte polarization ultimately lead to the delivery of mRNAs to specific regions of the oocyte, which causes a symmetry break of gene expression that defines head and tail of the future embryo (St Johnston and Nüsslein-Volhard, 1992). The main posterior determinant is *oskar* mRNA, which is delivered to the posterior by two distinct processes – directed transport and cytoplasmic streaming (Lu et al., 2018). Delivery by kinesin-dependent directed transport occurs during stages 7–9 of oogenesis. Because plus end–directed kinesin transport is stochastic and insensitive to Par-1, the microtubule cytoskeleton must be polarized so that microtubule plus–ends preferentially accumulate at the posterior. This is achieved through Par-1 dependent inhibition of microtubule nucleation at the posterior (Doerflinger et al., 2010; Nashchekin et al., 2016; Parton et al., 2011). Our results show that cell contact with PFCs inhibits Bazooka accumulation and retains Par-1 localization at the posterior of the oocyte. Therefore, PFCs have the crucial role in maintaining polarization of the microtubule cytoskeleton during stages when *oskar* mRNA is delivered to the posterior by microtubule-mediated directed transport. However, we also demonstrate that *oskar* accumulation at the posterior depends on Par-1 presence, and loss of posterior *oskar* occurs prior to changes in microtubule nucleation. At stage 10B, the mechanism of directed transport is substituted by cytoplasmic streaming, which delivers mRNA particles in bulk to the posterior where *oskar* is anchored by myosin V (Lu et al., 2018). Interestingly, this is the stage at which we observe loss of contact between the PFCs and the oocyte, as well as the loss of Par polarity. However, since polarization of the microtubule network is not necessary for cytoplasmic streaming, this does not compromise *oskar* localization during late oogenesis. Instead of maintaining tight contact between PFCs and the oocyte, molecular components building the eggshell can now be deposited into the intercellular space by the follicle cells to prepare for egg maturation (Cavaliere et al., 2008).

In our current understanding of Par protein interaction and Par domain formation (reviewed in Hoege and Hyman, 2013), the affinity of the posterior Par-1 for binding sites at the plasma membrane is modulated by phosphorylation activity of anterior Par complex member aPKC. Par-1 can phosphorylate anterior Par complex member Bazooka, which modulates the affinity of the anterior Par complex to bind to the membrane or binding sites thereon. Because Par proteins exhibit diffusive properties at the membrane (Goehring et al., 2011a), they not only accumulate by recruitment from the cytoplasm, but also move laterally on the membrane and form reciprocal domains, where they remain unphosphorylated and have high binding affinity. The antagonistic effect between anterior and posterior Pars is likely occurring everywhere in the cell but is strongest where these proteins are most concentrated – at the plasma membrane. This model can recapitulate Par domain formation, but it requires a trigger or initial symmetry break that allows the reaction-diffusion network to converge towards stable domain formation. In *C. elegans* the symmetry break is likely linked to the position of the centrosome provided by the sperm (Bienkowska and Cowan, 2012; Goldstein and Hird, 1996). A biochemical signal from the centrosome was proposed to locally inhibit actomyosin contractility and cause a local asymmetry in cortical flow (Munro et al., 2004). Since anterior Pars are embedded within the cortex, they are displaced from the posterior by advection allowing Par-1 (and Par-2) to accumulate from the cytoplasm (Goehring et al., 2011b; Munro et al., 2004). While the inhibitory signal (cascade) emerging from the centrosome is yet to be fully resolved (see Gan & Montegi, 2021 for a review), it is important to recognize in this model that signal transmission is achieved by a mechanical process that removes the anterior Par complex from what becomes the posterior domain. We show in the present study that, in the *Drosophila* oocyte, Bazooka exclusion – or inhibition of its accumulation – at the posterior involves the mechanical transduction of posterior follicle cell contact. This contact could either change the membrane composition, binding sites thereon, or the hydrodynamic events close by, leading to locally distinct physical and chemical circumstances for anterior Par proteins not to concentrate.

Both accumulation of Bazooka and delocalization of Par-1 following ablation of PFCs are restricted to the membrane that was in direct contact with ablated follicle cells. We conclude that the remaining PFCs continue transducing the signal that excludes Bazooka and retains Par-1. This agrees with previously reported work studying mosaic mutants in components of signaling pathways that are necessary for differentiation of PFCs (Frydman and Spradling, 2001; Poulton and Deng, 2006; Xi et al., 2003). In sum, these studies showed that *oskar* mRNA and Staufen protein correctly localize to the membrane facing WT follicle cells, while their localization is lost at the membrane facing neighboring cells that did not adopt posterior fate. This observation argues against the idea that the polarizing signal is a diffusible molecule within the extracellular space.

What distinguishes the cell-cell contact to PFCs from that of lateral follicle cells? Recent screening found that components of extracellular matrix (ECM) and ECM-associated proteins are upregulated in PFCs (Wittes and Schüpbach, 2019). Growing body of evidence suggests that there is a crosstalk between cell-ECM and cell-cell adhesion (reviewed in Mui et al., 2016). For example, integrins have been reported to increase the strength of cadherin adhesions (Martinez-Rico et al., 2010). Therefore, it is possible that PFCs-ECM interaction affects the adhesion between the PFCs and the oocyte. Alternatively, differential expression of eggshell genes occurs in the different subtypes of follicle cells. The eggshell is composed of five distinct layers, for which the molecular components are secreted by the follicle cells. The first layer is the vitelline membrane, which starts to be deposited at stage 9, at the time when Bazooka is excluded from the posterior (Cavaliere et al., 2008). There are four vitelline membrane genes, one of which, *VM32E*, is expressed in the main body follicle cells but not in the PFCs (Gargiulo et al., 1991). At stage 10, VM32E protein is found at the interface between main body follicle cells and the oocyte, but not at the posterior (Andrenacci et al., 2001). Interestingly, the protein spreads to the posterior by stage 11, which is the time when the connection between the PFCs and the oocyte is lost. Therefore, it is possible that VM32E is involved in separating follicle cells from the oocyte. Alternatively, the loss of contact between the PFCs and the oocyte could be the consequence of deposition of the second layer of eggshell, a vax layer, which starts at late stage 10 (Cavaliere et al., 2008).

Whatever the mechanism of keeping the oocyte and PFCs in close contact is, it seems to be important to maintain Bazooka exclusion. Interestingly, the first anterior-posterior polarization event - positioning of the oocyte to the posterior of the germline cyst at stage 1– is facilitated by cadherin mediated adhesion between follicle cells and the oocyte (Godt and Tepass, 1998; González-Reyes and St Johnston, 1998). In addition, the Par network becomes transiently polarized around this stage, with Par-1 at the posterior, and Bazooka at the anterior of the oocyte (Vaccari and Ephrussi, 2002). Based on this, it has been suggested that follicle cells are also involved in this first polarization of the oocyte (Roth and Lynch, 2009).

A change in adhesion between PFCs and the oocyte has been proposed as a mechanism by which the polarizing signal could be transferred from the PFCs to the oocyte at stage 7 (Poulton and Deng, 2007). How could the adhesion between follicle cells and the oocyte translate into oocyte polarization? Signals derived from cell-cell contact are regulators of polarization in many cells and contexts (reviewed in Ebnet et al., 2018). The most obvious downstream target of adhesion between the PFCs and the oocyte is the actin cytoskeleton. Cell adhesion modulates actin organization and dynamics (reviewed in Bachir et al., 2017). On the other hand, intact actin cytoskeleton is required both for posterior enrichment of Par-1 and exclusion of Bazooka (Doerflinger et al., 2006; Jouette et al., 2019). Recently, myosin activation at the posterior of the oocyte has been identified as the first known signal of oocyte polarity following the signal from PFCs. A continuous local dephosphorylation of myosin regulatory light chain (MRLC) is necessary for Par-1 localization (Doerflinger et al., 2022) and if would be interesting to understand the connection between PFC contact signaling and MRCL phosphorylation state. It has been proposed that activated myosin increases tension at the oocyte posterior, which might be necessary for recruitment of Par-1. cadherin has been reported to promote recruitment and activation of myosin in epithelia (Shewan et al., 2005). Therefore, it is possible that adhesion between the oocyte and PFCs causes specific activation of myosin at the posterior. Alternatively, the polarizing cue could be transferred from PFCs to the oocyte through a trafficking dependent process. Endocytosis is elevated at the posterior of the oocyte, and this has been linked to posterior localization of *oskar* (Tanaka and Nakamura, 2008; Vanzo et al., 2007). Additionally, Bazooka is not excluded from the posterior following either knockdown of Rab-5 or expression of a dominant negative form of Rab-5 in the oocyte. On the contrary, overexpression of the PIP5Kinase Skittles (SKTL), which produces phosphoinositide PI(4,5)P2, bypasses the need for Par-1 to have Bazooka excluded from the posterior. PI(4,5)P2 plays a role in the first steps of endocytosis, suggesting that SKTL overexpression rescues Bazooka exclusion in *par-1* mutants by increasing endocytosis (Jouette et al., 2019). The trafficking could work either through direct delivery of a polarizing signal or by a passive process, for example by remodeling the posterior membrane or through membrane flows (Gerganova et al., 2021). Thus, future work should focus on the biophysics and molecular key players governing plasma membrane dynamics specifically at the oocyte–PFC interface.

## Materials and Methods

### Fly stocks and husbandry

Fly husbandry was conducted in agreement with Portuguese National Regulations on Animal Welfare. Fly lines used in this study:

| Genotype | Origin / Gift | Reference |
| --- | --- | --- |
| w; +; Jupiter::mCherry | Daniel St Johnston Lab | Lowe et al., 2014 |
| w; +; Jupiter::GFP | Bloomington Drosophila Stock Center | #6836 |
| w; UASp>GFP::Par-1(N1S)/CyO;+ | Daniel St Johnston Lab | Huynh et al., 2001 |
| w; UASp>Bazooka::mGFP/CyO;+ | Daniel St Johnston Lab | Benton and St Johnston, 2003b |
| w; UASp>Bazooka::mCherry/CyO;+ | Bloomington Drosophila Stock Center | #65844 |
| w; GFP::aPKC; + | Eurico Morais-de-Sá Lab | Chen et al., 2018 |
| w; mat-α4>Gal4; + | Bloomington Drosophila Stock Center | #7062 |
| w; +; mat-α4>Gal4 | Bloomington Drosophila Stock Center | #7063 |
| w; +; osk>osk::MS2(10x)/TM3, Sb | Anne Ephrussi Lab | Zimyanin et al., 2008 |
| w; hsp83>MCP::mCherry/CyO; + | Anne Ephrussi Lab | Gáspár et al., 2017; Weil et al., 2006 |

The UASp>GFP::Par-1(N1S) isoform rescues the *par-1* mutant (Doerflinger et al., 2006). The rescuing activity of the UASp>Bazooka::mGFP transgene was demonstrated by Benton and St Johnston, 2003b. To express UASp-transgenes, flies were crossed with mat-α4>Gal4 driver. The cross was kept at 25°C for 3-4 days, after which parents were removed from the vial, and the vial was transferred to 18°C for the remaining time of the development. Viability and fertility tests were performed for all lines. Female offspring of the desired genotype were collected in vials with 3-4 males and supplied with fresh yeast. The flies were kept in vials at 18°C for 3-4 days before dissection. Detailed genotypes of flies used in the experiments were:

w*, mata4>Gal4 / UASp>Bazooka::mGFP ; Jupiter:mCherry/Jupiter::mCherry (Fig. 1–3, Suppl. Fig. 1A–B and Suppl. Fig. 2)

w*; UASp>GFP::Par-1(N1S)/+ ; mata4>Gal4/Bazooka::mCherry (Suppl. Fig. 1C–E; Fig. 4, Suppl. Fig. 3D–F)

w*; mata4>Gal4 / UASp>GFP::Par-1(N1S) ; Jupiter:mCherry/Jupiter::mCherry (Suppl. Fig. 3A–C)

w*; UASp>GFP::Par-1(N1S)/hsp83>MCP::mCherry; mata4>Gal4 /osk>osk::MS2(10x) (Fig. 5A–C)

w*; +/hsp83>MCP::mCherry; Jupiter::GFP/osk>osk::MS2(10x) (Fig,.5D–E)

w*; GFP::aPKC / UASp>Bazooka::mCherry; + / mata4>Gal4 (Fig. 6)

### Cleaning of coverslips and sample preparation

22×22 mm No. 1.5 coverslip (Marienfeld) were placed in a ceramic rack, placed into a beaker with NaOH (3M) and sonicated for 10 min. The rack was dipped-and-drained in a beaker with MilliQ water, transferred to clean MilliQ water and sonicated for 10 min. Finally, the rack was transferred into a new beaker with clean MilliQ water and sonicated for another 10 min. Coverslips were spin-dried and stored in a clean rack and sealed container until final use.

For experiments shown in Fig. 1–3, ovaries were dissected in a drop of halocarbon oil (Voltalef 10S, Arkema) placed on a clean coverslip, using tweezers to separate individual germaria. For experiments shown in Fig. 4–6, ovaries were dissected in a drop of Schneider’s medium supplemented with 10% FBS and 200 μg/mL insulin. Dissected ovaries were incubated 2×30s in 20 μL of supplemented Schneider’s medium. Finally, ovaries were transferred to a drop of supplemented Schneider’s medium on a clean coverslip next to a drop of halocarbon oil and individual germaria were pulled into the oil. This latter protocol improved the sample lifetime and allowed for longer time-lapse imaging.

### Microscopy

Imaging was performed on a Nikon Eclipse Ti-E microscope equipped with a Yokogawa CSU-W Spinning Disk confocal scanner and a piezoelectric stage (737.2SL, Physik Instrumente), using 40x water immersion objective and 488 nm and 561 nm laser lines for excitation of GFP and mCherry, respectively. Images were acquired at 73 focal planes with 0.5 μm z-spacing. x-y pixel size was 162 nm. An Andor iXon3 888 EMCCD camera was used for time-lapse acquisitions, while an Andor Zyla 4.2 sCMOS camera, which offers 2x wider field of view, was used for snapshot images of whole egg chambers for analyses as shown in Fig. 1 and Suppl. Fig. 1. During time-lapse microscopy, images were acquired every 30 s (in experiments show in Fig. 2–3, Suppl Fig. 2, and Suppl. Fig. 3A–C) or every 60 s (in experiments show in Fig. 4 and Suppl. Fig. 3D–F) for 60–90 minutes. To assess Par protein dynamics at longer timescales (Fig. 5 and Fig. 6), images were acquired every 5 min for at least 150 minutes.

### Laser ablation

The laser ablation system used in this study was described elsewhere (De-Carvalho et al., 2022). Follicle cells were ablated using circular ablation (60 px diameter, 5 px step size). Ablation was performed several times while moving manually in z direction to UV expose the entire depth of the cells. In samples which were not dissected in Schneider’s medium, ablation was performed 25 times using 50% laser power. When sample was dissected in Schneider’s medium laser ablation became more effective, presumably because the egg chambers are surrounded by a thin layer of aqueous solution. Therefore, ablation in these samples was performed 5 times using 20% laser power. To control for unspecific effects of laser ablation, the ooplasm was ablated using identical power settings.

### Fluorescence recovery after photobleaching (FRAP)

Par1::GFP at the oocyte membrane was bleached along ∼100 px long line using the laser used for the ablation (2% laser power, 1 px step size). Bleaching was performed in 10 z-planes with 0.5 μm spacing between planes. Images were acquired every 15 s for 15 min. At least two images before photobleaching were acquired.

### Image processing

Image deconvolution was done in Huygens (Scientific Volume Imaging) using a theoretical point spread function (PSF), 40 iterations, a signal-to-noise ratio of 8 and automatic brick layout. The background was estimated by measuring the signal in areas where there was no egg chamber. Deconvolution was done separately for each channel. Images shown in the figures were made by calculating the sum of 6 z-slices in Fiji/ImageJ (National Institutes of Health; (Schindelin et al., 2012). Final figures were assembled in Illustrator (Adobe).

### Image analysis

All measurements were performed in Fiji/ImageJ on a sum of 6 z-slices. All measurements were performed on raw images, except for the intensity profiles shown in Fig. 1A-C and Fig. 2D, which were done on deconvolved images. The profiles were measured by drawing a 10 px wide line from the oocyte to the follicle cells and measuring the mean intensity across the line in all three channels (Bazooka::GFP, Jupiter::mCherry and polarized transmission light).

Profiles of signal intensities at the posterior membrane of the oocytes were calculated by drawing a 10 px wide segmented line across the oocyte posterior cortex. Values of intensities for each frame were normalized using min-max normalization. The zero position on the x-axis represents the reference point, i.e. the polar cells or the ablation spot.

To calculate the posterior to lateral intensity ratio, a 10 px wide and ∼50 μm long line was drawn at the posterior and lateral membrane cortex of the oocyte. Mean intensity across this line was calculated and the mean value of the background signal was subtracted from this value. The background signal was measured by drawing a 10 px wide and ∼50 μm long line in an area of the image where there was no egg chamber.

To measure the change in intensity over time, a 10 px wide segmented line was drawn, covering only the membrane that was previously in contact with ablated follicle cells. The line was manually moved if necessary due to x-y drift of the sample. Mean value of intensity across this line was measured in all time frames. Background signal was measured in the ooplasm next to the posterior membrane of the oocyte, using a 10 px wide and ∼50 μm long line. Intensity shown in graphs was calculated by subtracting the mean value of the background signal from the mean intensity at the membrane. To produce the heatmaps, the profile of intensity across the line was obtained, and mean background signal was subtracted. Since the length of the ablation region was different in different experiments, the values of pixel intensities were binned into 30 bins. The final heatmaps show the intensity in 30 bins averaged over several experiments.

Mean fluorescence intensity of the photobleached region was measured by drawing a 10 px wide line over the region. To correct for photobleaching, the mean intensity of reference signal was measured inside the ooplasm. The background signal, measured outside of the egg chamber, was subtracted from the signal measured in both bleached and reference region. Normalized intensity was calculated as:

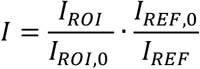

Where *I*_*ROI*_ and *I*_*REF*_ are background subtracted intensity at the bleached region and reference region, respectively. *I*_*ROI*,0_ and *I*_*REF*,0_ are intensities before bleaching.

The normalized intensity was fitted to single exponential equations of the form:

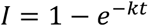

Finally, the half-time of recovery was calculated as: 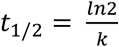

Signal decays in Fig. 5 were fitted to a single exponential function with time delay *τ* as fitting parameter:

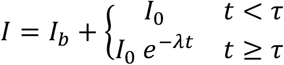

where *I*_*b*_ is the baseline intensity and *I*_0_ is the signal amplitude decaying at rate *λ*.

Statistical analysis, curve fitting and plotting were done using Python. Information on sample size, statistical tests and p-values are shown in figures or mentioned in figure captions.

## Acknowledgements

We thank members of the Telley lab for fruitful discussions. We acknowledge I. Gáspár and A. Ephrussi for suggestions on sample preparation. We thank the staff of the Fly Facility, the Advanced Imaging Facility (AIF) and the Technical Support Service at the Instituto Gulbenkian de Ciência (IGC). We thank Anne Ephrussi (EMBL Heidelberg), Daniel St. Johnston (The Gurdon Institute) and Eurico Morais-de-Sá (I3S Porto) for fly line donations, and The Bloomington Drosophila Stock Center for their services.

## Funding

We acknowledge financial support from: Human Frontiers Science Program (HFSP) awarded to I.A.T. and supporting A.M. and J.C. (RGY0083/2016); the Fundação para Ciência e a Tecnologia (FCT) supporting I.A.T. (Investigador FCT IF/00082/2013) and A.M. (2020.04666.BD); Fundação Calouste Gulbenkian (FCG) and LISBOA-01-0145-FEDER-007654 supporting IGC’s core operation; LISBOA-01-7460145-FEDER-022170 (Congento) supporting the Fly Facility; PPBI-POCI-01-0145-FEDER-022122 supporting the AIF, all co-financed by FCT (Portugal) and Lisboa Regional Operational Program (Lisboa2020) under the PORTUGAL2020 Partnership Agreement (European Regional Development Fund).

## Author contributions

A.M. J.C., I.A.T. conceived the study; A.M. and J.C. established and validated the fly lines; A.M. designed and carried out all the experiments; A.M. and I.A.T. analyzed the data; I.A.T. designed and assembled the microscopy, micromanipulation, and laser ablation platform; A.M. wrote the draft manuscript guided by I.A.T.; all authors edited the final manuscript; I.A.T. supervised the project.

## Competing interests

The authors have no competing financial or non-financial interests.

## Supplementary Figures

**Suppl. Figure 1:**
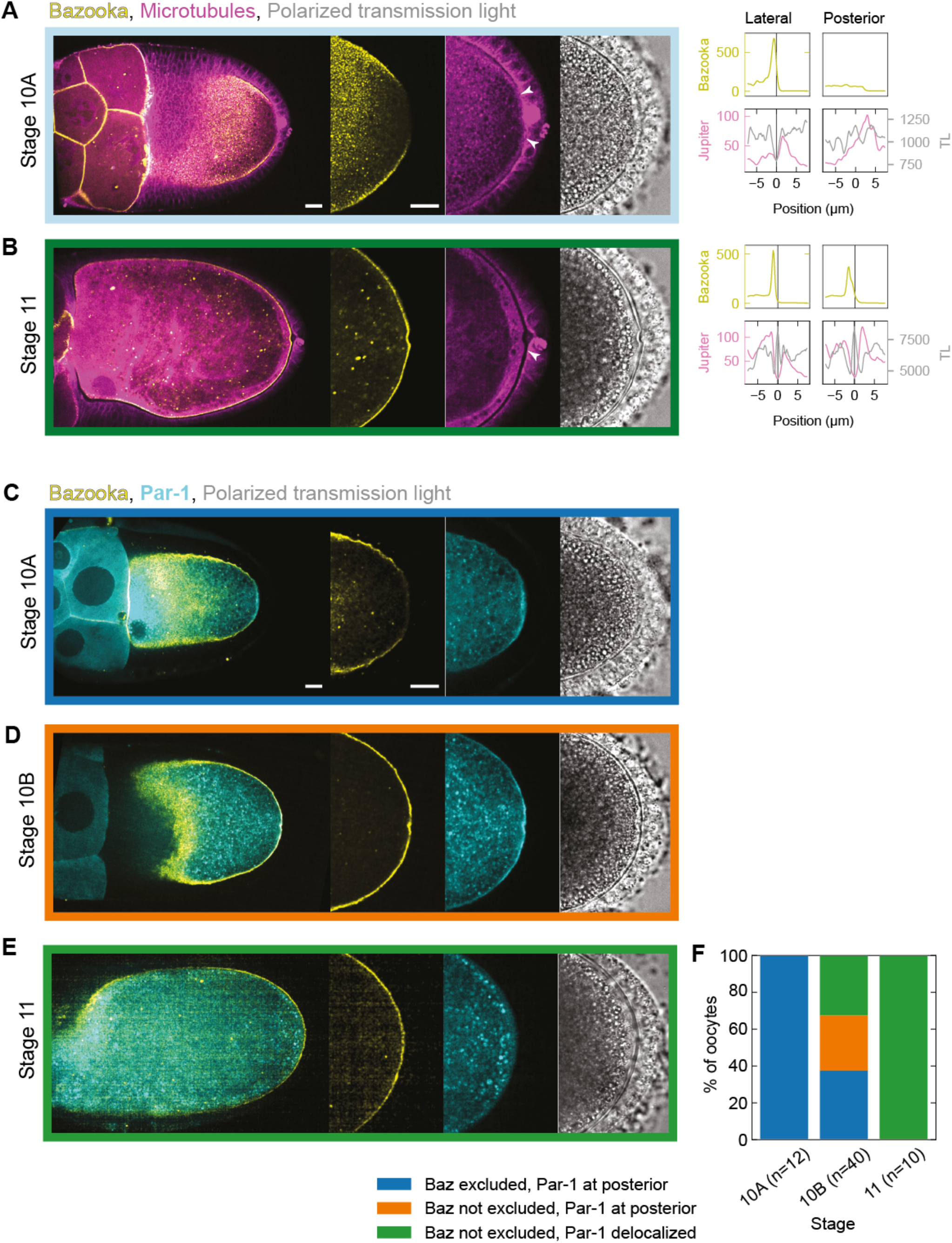
Localization of Bazooka and Par-1 during stages 10 and 11 of oogenesis: On the left are still images of stage 10A **(A)** and 11 **(B)** egg chambers expressing Bazooka::GFP (yellow) and the microtubule reporter Jupiter::mCherry (magenta) next to a transmission light micrograph (grey). The right graphs are intensity profiles of Bazooka::GFP (yellow), Jupiter::mCherry (magenta) and transmission light (grey) signal along a straight line crossing cell boundaries from the oocyte to either lateral or posterior follicle cells. The x-axis origin and the vertical line represent the local minimum of the Jupiter::mCherry (magenta) signal, which we interpret as extended intercellular space (gap) between the oocyte and the follicle cells. If the intercellular space is not clearly discernible, the position 0 μm represents the midpoint of the line. **(A)** At stage 10A, the gap is visible between the oocyte and the lateral follicle cells, but not between the oocyte and the posterior follicle cells. Accumulation of Bazooka correlates with the existence of the gap. **(B)** At stage 11, both the gap and Bazooka accumulation are visible at the posterior. **(C–E)** Stage 10A, 10B and 11 egg chambers expressing Bazooka::mCherry (yellow) and Par-1::GFP (cyan) in the germline. At stage 10A, the mutual exclusion zone between Bazooka and Par-1 exists at the posterior. During stage 10B Bazooka accumulates to the posterior before delocalization of Par-1. At stage 11, both exclusion of Bazooka and accumulation of Par-1 at the posterior are lost. **(F)** Distribution of egg chambers showing different posterior Par protein localization patterns. The colors in the plot correspond to the color of the frame surrounding the images in panels C–E. Note that Par-1 delocalization never precedes accumulation of Bazooka. Scale bars represent 20 μm, *n* is the number of egg chambers.

**Suppl. Figure 2:**
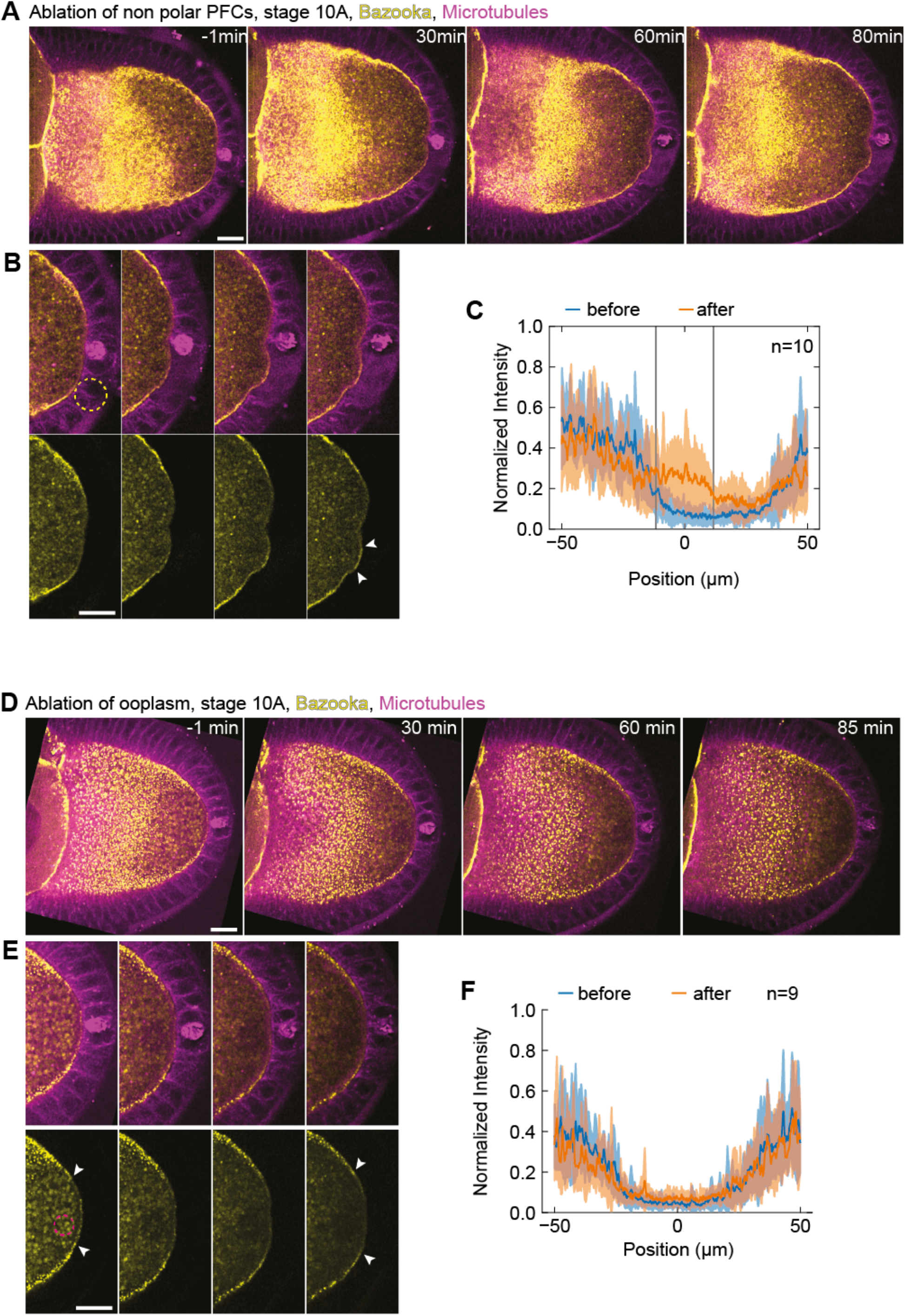
Bazooka localizes to the posterior following ablation of PFCs: **(A)** Two-color time-lapse images of an egg chamber expressing Bazooka::GFP in the germline (yellow) and the microtubule marker Jupiter::mCherry (magenta) before (-1 min) and after ablation of PFCs. **(B)** Zoom-in of the posterior of the egg chamber shown in panel A. Top: merged channels, bottom: Bazooka::GFP. Dashed circle in the first image of the top panel represents the ablation spot. Note that polar cells (pair of cells with high Jupiter signal) are not ablated. After 80 min, the accumulation of Bazooka::GFP is visible at the oocyte membrane facing the ablated cells (arrowheads in last image). **(C)** Average intensity profile of Bazooka::GFP signal at the posterior of the oocyte before (blue) and after (orange) ablation. The shaded region represents the standard deviation. The x-axis origin represents the position of the ablated region. Vertical lines denote the average width of the ablated region. Note that the signal increases markedly only within the ablated region. **(D)** Two-color time-lapse images of an egg chamber expressing Bazooka::GFP in the germline (yellow) and microtubule marker Jupiter::mCherry (magenta) before (-1 min) and after ablation within the ooplasm. **(E)** Zoom-in of the posterior of the egg chamber shown in panel D. Top: merged channels, bottom: Bazooka::GFP. Dashed circle in the first image of the bottom panel shows the position of the ablation spot. Bazooka remains excluded from the posterior after ablation (arrowheads). **(F)** Average intensity profile of Bazooka::GFP signal at the posterior of the oocyte before (blue) and after (orange) ablation within the ooplasm. The shaded region designates the standard deviation. The x-axis origin represents the position of polar cells. For all panels the scale bar represents 20 μm, and *n* is the number of experiments.

**Suppl. Figure 3.**
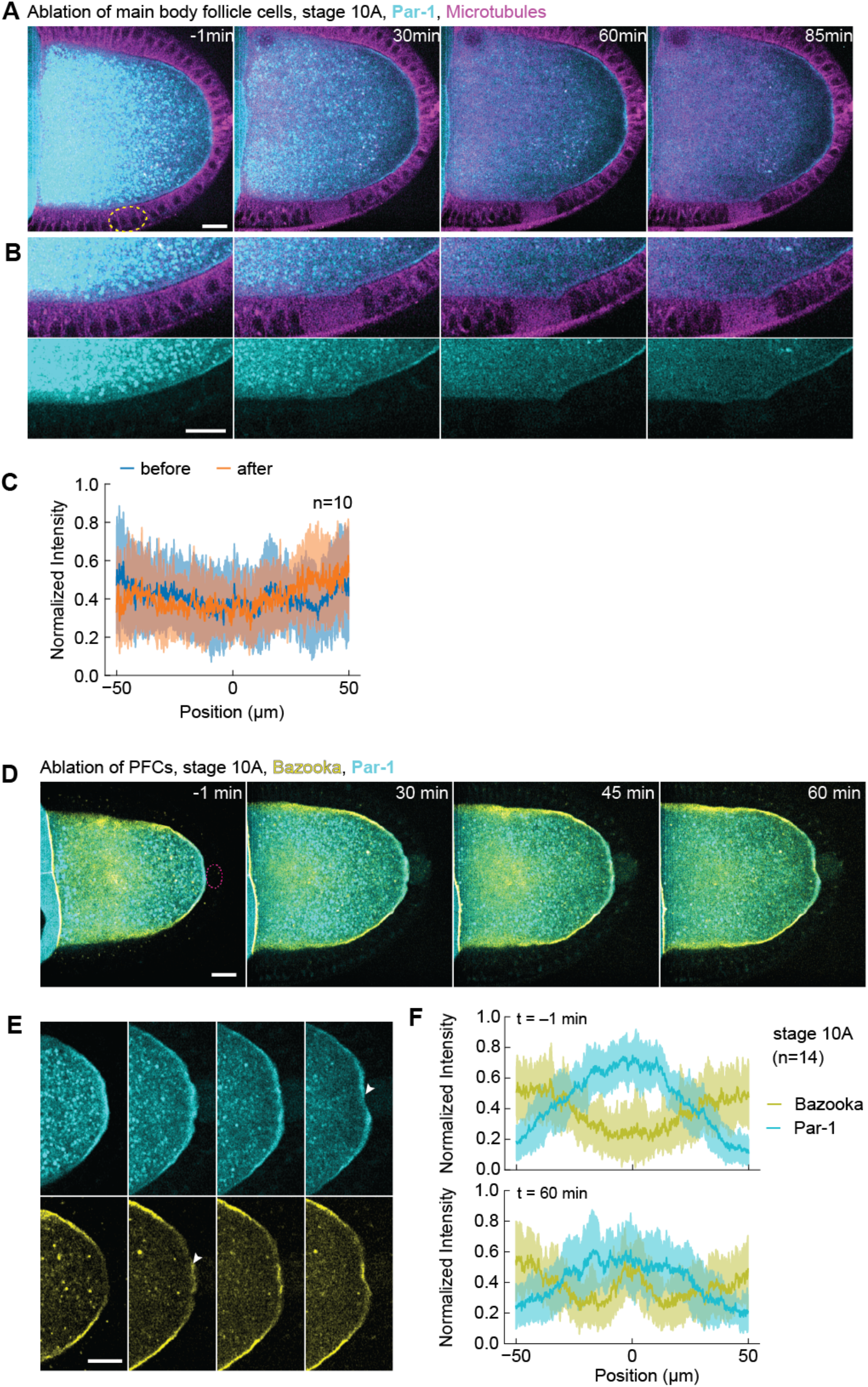
Par-1 loss from the posterior following PFC ablation is slower in stage 10A oocytes. **(A)** Two-colour time-lapse images of a control experiment on an egg chamber expressing Par-1::GFP in the germline (cyan) and Jupiter::mCherry (magenta) before (-1 min) and after ablation of main body follicle cells (yellow dashed circle). **(B)** Zoom-in of the lateral side of the egg chamber shown in panel A. Top: merged channels, bottom: Par-1::GFP. No change in Par-1 localization is observed. **(C)** Average intensity profile of Par-1::GFP signal around the ablated region before (blue) and after (orange) ablation of the main body follicle cells. The shaded region designates the standard deviation. The x-axis origin represents the center of the ablated region. **(D)** Two-color time-lapse images of an egg chamber at stage 10A expressing Par1::GFP (cyan) and Bazooka::mCherry (yellow) in the germline, before (-1 min) and after ablation of PFCs (red dashed circle). Scale bar, 20 μm **(E)** Zoom into the posterior of the egg chamber shown in panel A. Top: Par1::GFP, bottom: Bazooka::mCherry. Following ablation, Bazooka accumulates to the posterior, while Par-1 marginally delocalizes (arrowheads). Scale bar, 20 μm. **(F)** Average intensity profile of Par-1::GFP (cyan) and Bazooka::mCherry (yellow) signal at the posterior of stage 10A oocytes before (top graph) and one hour after ablation of PFCs (bottom graph). The shaded region designates the standard deviation. The x-axis origin represents the position of the polar cells (top) or center of ablation spot (bottom). Note that Par-1 signal does not decrease much.

## Supplementary Video Legends

**Video 1**. Sum of 6 z-slices from a time-lapse movie of stage 10A egg chamber expressing Bazooka::GFP (yellow) and microtubule marker Jupiter::mCherry (magenta) following detachment of PFCs from the oocyte. Left: polarized transmission light, right: florescence. Frame rate is 10 frames/s. In support of Fig. 2.

**Video 2**. Sum of 6 z-slices from a time-lapse movie of stage 10A egg chamber expressing Bazooka::GFP (yellow) and microtubule marker Jupiter::mCherry (magenta). PFCs were ablated at time 0. Frame rate is 10 frames/s. In support of Fig. 3.

**Video 3**. Sum of 6 z-slices from a time-lapse movie of stage 10A egg chamber expressing Bazooka::GFP (yellow) and microtubule marker Jupiter::mCherry (magenta). Ooplasm was ablated at time 0. Frame rate is 10 frames/s. In support of Suppl. Fig. 2.

**Video 4**. Sum of 6 z-slices from a time-lapse movie of stage 9 egg chamber expressing Par1::GFP (cyan) and Bazooka::mCherry (yellow). PFCs were ablated at time 0. Frame rate is 10 frames/s. In support of Fig. 4.

**Video 5**. Sum of 6 z-slices from a time-lapse movie of stage 10A egg chamber expressing Par1::GFP (cyan) and Bazooka::mCherry (yellow). PFCs were ablated at time 0. Frame rate is 10 frames/s. In support of Suppl. Fig. 3.

**Video 6**. Sum of 6 z-slices from a time-lapse movie of stage 9 egg chamber expressing Par1::GFP (cyan) and *oskar::MS2*-mCherry (magenta). PFCs were ablated at time 0. Frame rate is 5 frames/s. In support of Fig. 5.

**Video 7**. Sum of 6 z-slices from a time-lapse movie of stage 9 egg chamber expressing microtubule marker Jupiter::GFP (green) and *oskar::MS2*-mCherry (magenta). PFCs were ablated at time 0. Frame rate is 5 frames/s. In support of Fig. 5.

**Video 8**. Sum of 6 z-slices from a time-lapse movie of egg chamber expressing Bazooka::mCherry (magenta) in the germline and aPKC::GFP (green). PFCs were ablated at time 0. Frame rate is 5 frames/s. In support of Fig. 6

## Notes

### Competing Interest Statement

The authors have declared no competing interest.

### Summary of Updates

In the revised manuscript we have added additional experimental data showing that downstream of Par polarity in the oocyte, posterior follicle cell contact is also required to maintain oskar mRNA localisation at the posterior.

## References

Andrenacci, D., Cernilogar, F.M., Taddei, C., Rotoli, D., Cavaliere, V., Graziani, F., and Gargiulo, G. (2001). Specific domains drive VM32E protein distribution and integration in Drosophila eggshell layers. J. Cell Sci. 114, 2819–2829.

Bachir, A.I., Horwitz, A.R., Nelson, W.J., and Bianchini, J.M. (2017). Actin-based adhesion modules mediate cell interactions with the extracellular matrix and neighboring cells. Cold Spring Harb. Perspect. Biol. 9:a023234, 1–19.

Benton, R., and St Johnston, D. (2003a). Drosophila PAR-1 and 14-3-3 Inhibit Bazooka / PAR-3 to Establish Complementary Cortical Domains in Polarized Cells. Cell 115, 691–704.

Benton, R., and St Johnston, D. (2003b). A Conserved Oligomerization Domain in Drosophila

Bazooka/PAR-3 Is Important for Apical Localization and Epithelial Polarity. Curr. Biol. 13, 1330–1334.

Bienkowska, D., and Cowan, C.R. (2012). Centrosomes can initiate a polarity axis from any position within one-cell C. Elegans embryos. Curr. Biol. 22, 583–589.

Cavaliere, V., Bernardi, F., Romani, P., Duchi, S., and Gargiulo, G. (2008). Building up the Drosophila eggshell: First of all the eggshell genes must be transcribed. Dev. Dyn. 237, 2061–2072.

Chen, J., Sayadian, A.C., Lowe, N., Lovegrove, H.E., and St Johnston, D. (2018). An alternative mode of epithelial polarity in the Drosophila midgut. PLoS Biol. 16, 1–24.

De-Carvalho, J., Tlili, S., Hufnagel, L., Saunders, T.E., and Telley, I.A. (2022). Aster repulsion drives short-ranged ordering in the Drosophila syncytial blastoderm. Development 149, 1–13.

Doerflinger, H., Benton, R., Torres, I.L., Zwart, M.F., and St Johnston, D. (2006). Drosophila Anterior-Posterior Polarity Requires Actin-Dependent PAR-1 Recruitment to the Oocyte Posterior. Curr. Biol. 16, 1090–1095.

Doerflinger, H., Vogt, N., Torres, I.L., Mirouse, V., Koch, I., Nüsslein-volhard, C., and St Johnston, D. (2010). Bazooka is required for polarisation of the Drosophila anterior-posterior axis. Development 137, 1765–1773.

Doerflinger, H., Zimyanin, V., and St Johnston, D. (2022). The Drosophila anterior-posterior axis is polarized by asymmetric myosin activation. Curr. Biol. 32, 374-385.e4.

Ebnet, K., Kummer, D., Steinbacher, T., Singh, A., Nakayama, M., and Matis, M. (2018). Regulation of cell polarity by cell adhesion receptors. Semin. Cell Dev. Biol. 81, 2–12.

Frydman, H.M., and Spradling, A.C. (2001). The receptor-like tyrosine phosphatase Lar is required for epithelial planar polarity and for axis determination within Drosophila ovarian follicles. Development 128, 3209–3220.

Gargiulo, G., Gigliotti, S., Malva, C., and Graziani, F. (1991). Cellular specificity of expression and regulation of Drosophila vitelline membrane protein 32E gene in the follicular epithelium: identification of cis-acting elements. Mech. Dev. 35, 193–203.

Gáspár, I., Sysoev, V., Komissarov, A., and Ephrussi, A. (2017). An RNA-binding atypical tropomyosin recruits kinesin-1 dynamically to oskar mRNPs. EMBO J. 36, 319–333.

Gerganova, V., Lamas, I., Rutkowski, D.M., Vještica, A., Castro, D.G., Vincenzetti, V., Vavylonis, D., and Martin, S.G. (2021). Cell patterning by secretion-induced plasma membrane flows. Sci. Adv. 7, 1–19.

Godt, D., and Tepass, U. (1998). Drosophila oocyte localization is mediated by differential cadherin-based adhesion. Nature 395, 387–391.

Goehring, N.W., Hoege, C., Grill, S.W., and Hyman, A.A. (2011a). PAR proteins diffuse freely across the anterior–posterior boundary in polarized. J. Cell Biol. 193, 583–594.

Goehring, N.W., Khuc Trong, P., Bois, J.S., Chowdhury, D., Nicola, E.M., Hyman, A.A., and Grill, S.W. (2011b). Polarization of PAR Proteins by Advective Triggering of a Pattern-Forming System. Science (80-.). 334, 1137–1141.

Goldstein, B., and Hird (1996). Specification of the anteroposterior axis in Caenorhabditis elegans. Development 122, 1467–1474.

González-Reyes, A., and St Johnston, D. (1994). Role of Oocyte Position in Establishment of Anterior-Posterior Polarity in Drosophila. Science (80-.). 266, 639–642.

González-Reyes, A., and St Johnston, D. (1998). The Drosophila AP axis is polarised by the cadherin-mediated positioning of the oocyte. Development 125, 3635–3644.

González-Reyes, A., Elliot, H., and St Johnston, D. (1995). Polarization of both major body axes in Drosophila by gurken-torpedo signalling. Nature 375, 654–658.

Hoege, C., and Hyman, A.A. (2013). Principles of PAR polarity in Caenorhabditis elegans embryos. Nat. Rev. Mol. Cell Biol. 14, 315–322.

Huynh, J.-R., Shulman, J.M., Benton, R., and St Johnston, D. (2001). PAR-1 is required for the maintenance of oocyte fate in Drosophila. Development 128, 1201–1209.

Jouette, J., Guichet, A., and Claret, S.B. (2019). Dynein-mediated transport and membrane trafficking control PAR3 polarised distribution. Elife 8:e402128, 1–28.

Lang, C.F., and Munro, E. (2017). The PAR proteins : from molecular circuits to dynamic self-stabilizing cell polarity. Development 144, 3405–3416.

Lowe, N., Rees, J.S., Roote, J., Ryder, E., Armean, I.M., Johnson, G., Drummond, E., Spriggs, H., Drummond, J., Magbanua, J.P., et al. (2014). Analysis of the expression patterns, Subcellular localisations and interaction partners of drosophila proteins using a pigp protein trap library. Development 141, 3994–4005.

Lu, W., Lakonishok, M., Serpinskaya, A.S., Kirchenbüechler, D., Ling, S.C., and Gelfand, V.I. (2018). Ooplasmic flow cooperates with transport and anchorage in Drosophila oocyte posterior determination. J. Cell Biol. 217, 3497–3511.

Lu, W., Lakonishok, M., Liu, R., Billington, N., Rich, A., Glotzer, M., Sellers, J.R., and Gelfand, V.I. (2020). Competition between kinesin-1 and myosin-V defines Drosophila posterior determination. Elife 9:e54216, 1–25.

Martinez-Rico, C., Pincet, F., Thiery, J.P., and Dufour, S. (2010). Integrins stimulate E-cadherin-mediated intercellular adhesion by regulating Src-kinase activation and actomyosin contractility. J. Cell Sci. 123, 712–722.

Motegi, F., and Seydoux, G. (2013). The PAR network: Redundancy and robustness in a symmetry-breaking system. Philos. Trans. R. Soc. B Biol. Sci. 368, 20130010.

Mui, K.L., Chen, C.S., and Assoian, R.K. (2016). The mechanical regulation of integrin-cadherin crosstalk organizes cells, signaling and forces. J. Cell Sci. 129, 1093–1100.

Munro, E., Nance, J., and Priess, J.R. (2004). Cortical flows powered by asymmetrical contraction transport PAR proteins to establish and maintain anterior-posterior polarity in the early C. elegans embryo. Dev. Cell 7, 413–424.

Nashchekin, D., Ribeiro Fernandes, A., and St Johnston, D. (2016). Patronin / Shot Cortical Foci Assemble the Noncentrosomal Microtubule Array that Specifies the Drosophila Anterior-Posterior Axis. Dev. Cell 38, 61–72.

Parton, R.M., Hamilton, R.S., Ball, G., Yang, L., Cullen, C.F., Lu, W., Ohkura, H., and Davis, I. (2011). A PAR-1–dependent orientation gradient of dynamic microtubules directs posterior cargo transport in the Drosophila oocyte. J. Cell Biol. 194, 121–135.

Poulton, J.S., and Deng, W.-M. (2006). Dystroglycan down-regulation links EGF receptor signaling and anterior-posterior polarity formation in the Drosophila oocyte. Proc. Natl. Acad. Sci. 103, 12775–12780.

Poulton, J.S., and Deng, W.M. (2007). Cell-cell communication and axis specification in the Drosophila oocyte. Dev. Biol. 311, 1–10.

Riechmann, V., and Ephrussi, A. (2001). Axis formation during Drosophila oogenesis. Curr. Opin. Genet. Dev. 11, 374–383.

Roth, S., and Lynch, J.A. (2009). Symmetry Breaking During Drosophila Oogenesis. Cold Spring Harb. Perspect. Biol. 1:a001891, 1–22.

Roth, S., Neuman-Silberberg, F.S., Barcelo, G., and Schüpbach, T. (1995). cornichon and the EGF Receptor Signaling Process Are Necessary for Both Anterior-Posterior and Dorsal-Ventral Pattern Formation in Drosophila. Cell 81, 967–978.

Schindelin, J., Arganda-Carreras, I., Frise, E., Kaynig, V., Longair, M., Pietzsch, T., Preibisch, S., Rueden, C., Saalfeld, S., Schmid, B., et al. (2012). Fiji: An open-source platform for biological-image analysis. Nat. Methods 9, 676–682.

Shewan, A.M., Maddugoda, M., Kraemer, A., Stehbens, S.J., Verma, S., Kovacs, E.M., and Yap, A.S. (2005). Myosin 2 Is a Key Rho Kinase Target Necessary for the Local Concentration of E-Cadherin at Cell–Cell Contacts. Mol. Biol. Cell 16, 4531–4542.

Shulman, J.M., Benton, R., and St Johnston, D. (2000). The Drosophila Homolog of C. elegans PAR-1 Organizes the Oocyte Cytoskeleton and Directs oskar mRNA Localization to the Posterior Pole. Cell 101, 377–388.

St Johnston, D., and Nüsslein-Volhard, C. (1992). The origin of pattern and polarity in the Drosophila embryo. Cell 68, 201–219.

Tanaka, T., and Nakamura, A. (2008). The endocytic pathway acts downstream of Oskar in Drosophila germ plasm assembly. Development 135, 1107–1117.

Tomancak, P., Piano, F., Riechmann, V., Gunsalus, K.C., Kemphues, K.J., and Ephrussi, A. (2000). A Drosophila melanogaster homologue of Caenorhabditis elegans par-1 acts at an early step in embryonic-axis formation. Nat. Cell Biol. 2, 458–460.

Vaccari, T., and Ephrussi, A. (2002). The fusome and microtubules enrich Par-1 in the oocyte, where it effects polarization in conjunction with Par-3, BicD, Egl, and Dynein. Curr. Biol. 12, 1524–1528.

Vanzo, N., Oprins, A., Xanthakis, D., Ephrussi, A., and Rabouille, C. (2007). Stimulation of Endocytosis and Actin Dynamics by Oskar Polarizes the Drosophila Oocyte. Dev. Cell 12, 543–555.

Weil, T.T., Forrest, K.M., and Gavis, E.R. (2006). Localization of bicoid mRNA in Late Oocytes Is Maintained by Continual Active Transport. Dev. Cell 11, 251–262.

Wittes, J., and Schüpbach, T. (2019). A gene expression screen in Drosophila melanogaster identifies novel JAK/STAT and EGFR targets during oogenesis. G3 Genes, Genomes, Genet. 9, 47–60.

Xi, R., McGregor, J.R., and Harrison, D.A. (2003). A gradient of JAK pathway activity patterns the anterior-posterior axis of the follicular epithelium. Dev. Cell 4, 167–177.

Zimyanin, V.L., Belaya, K., Pecreaux, J., Gilchrist, M.J., Clark, A., Davis, I., and St Johnston, D. (2008). In Vivo Imaging of oskar mRNA Transport Reveals the Mechanism of Posterior Localization. Cell 134, 843–853.

